# The concentric β-barrel hypothesis for amyloids: Models of soluble and transmembrane amyloid-β42 oligomers and channels composed of identical subunits and GM1 gangliosides

**DOI:** 10.64898/2026.03.19.711324

**Authors:** Stewart R. Durell, Yinon Shafrir, H. Robert Guy

**Author notes:** Contact information: H. Robert Guy Amyloid Research Consultants (ARC), 6510 Tahawash Street, Cochiti Lake, NM 87083, USA.

## Abstract

Soluble oligomers and transmembrane channels formed by the 42-residue variant of amyloid beta (Aβ42) play key roles in Alzheimer’s disease. Unfortunately, detailed structures of these assemblies have not been determined. Our group addresses this problem by developing atomic scale models. Previously we proposed that both soluble Aβ42 oligomers and transmembrane channels have symmetric concentric β-barrel structures. Here we expand this hypothesis to include GM1 gangliosides and sometimes cholesterol and lattice models of channel assemblies. The presence of GM1 gangliosides increases the toxicity of Aβ42, enhances its ability to penetrate liposome membranes, and facilitates interactions between adjacent liposomes. Although the conformations of numerous model assemblies vary, in these models the carboxyl group of GM1 always binds to side-chains of histidine 13 and/or histidine 14. Our soluble oligomer models are consistent with electron microscopy images of beaded annular protofibrils. Our models of membrane-bound assemblies are consistent with the following: freeze-fracture and atomic force microscopy images of Aβ42 in lipid bilayers, secondary structure results, the calcium hypothesis of Alzheimer’s Disease, effects of lithium depletion on AD, established β-barrel theory, and energetic criteria.

## Introduction

Aβ oligomers have multiple effects when they interact with lipids and membranes (see (Mrdenovic et al., 2022) for recent review); they can interact in solution with free lipids including GM1 ganglioside and cholesterol ((Sciacca et al., 2020), (Chakravorty et al., 2022), (Zhaliazka et al., 2023), (Hashemi et al., 2022)), they can exert detergent-like disruptions of membrane ((Bode et al., 2019)), they can extract lipids from membranes to form proteolipid assemblies ((Michikawa et al., 2001)), they are highly polymorphic ((Huang & Liu, 2020)), there are many variants ((Busch & Bufe, 2023)), they interact with numerous membrane proteins ((Wiatrak et al., 2021)), and in some situations they are disordered ((Coskuner-Weber et al., 2022)). Their effects vary depending upon types of lipids ((Budvytyte & Valincius, 2023)), the curvature of the membrane, the length of the Aβ peptide, the size of the oligomers, the source of the peptides, methods used to prepare the peptides and lipids, the ratio of lipids to peptide, and other cofactors. Also, Aβ oligomers affect numerous membranes; both synaptic from the outside ((John & Reddy, 2021)) and organelles (mitochondria, golgi apparatus, endosomes, and vesicles) from the inside ((Gallego Villarejo et al., 2022)). Uncertainty remains about which processes are functional and pathogenic ((Sulatskaya et al., 2021)).

Some would call efforts to determine precise three-dimensional structures of so-called “Intrinsically Disordered Protein’s (IDPs)” pre-fibril assemblies a fool’s errand. The validity of the IDP term for Aβ assemblies depends upon how it is defined and which form is being discussed. If taken to mean that the peptides have little or no regular secondary or tertiary structure, then it is applicable primarily to assemblies of only a few subunits in the aqueous phase and possibly to some Aβ variants. In contrast, fibrils composed of many Aβ subunits are highly ordered but polymorphic. Most Aβ fibrils have parallel β structure ((Stroud et al., 2012)), but others do not. Fibril stability can be attributed to the network of backbone hydrogen bonds and other energetically favorable interactions formed between adjacent subunits. These interactions are absent in monomers and reduced in small assemblies in water, so it is reasonable that they tend to be disordered. Numerous proteins and peptides are metamorphic, they exist on the boundary between complete disorder and highly ordered ((Kulkarni et al., 2018)).

Environmental factors such as peptide concentration, pH, temperature, post-translational modifications, time, and interactions with other molecules (e.g., lipids, fatty acids, membranes, other proteins, ions), can trigger such polymorphic and/or metamorphic proteins to transition among different topologies. Aβ assemblies exhibit these properties, they adopt antiparallel β-secondary structure forms as the number of subunits increases to form oligomers and protofibrils and when hydrophobic solvents, detergents, lipids, and/or membranes are present ((Cerf et al., 2009), (Kayed et al., 2009)). Thus, relatively small Aβ assemblies with a high content of regular secondary structure are still polymorphic.

Although it has long been known that Aβ forms channels in lipid bilayers ((Arispe et al., 1993)); Aβ42 is the only variant to form discrete channels in neurons ((Bode et al., 2017)). Here we focus only on Aβ42 because it appears to be the most toxic variant ((Fu et al., 2017)) and is the variant used to obtain freeze-fracture EM images that constrain our models ((Shafrir, Durell, Arispe, et al., 2010; Shafrir, Durell, Anishkin, et al., 2010)). Some of our current models include GM1 ganglioside and/or cholesterol, which increase the toxicity of Aβ42 assemblies and binding to membranes ((Zhang et al., 2022), (Fantini et al., 2020)). A few channel models include lithium which has been reported to inhibit onset of AD (Aron et al., 2025; Terao & Kodama, 2024). The structures of Aβ42 assemblies differ from those of the other three most common variants (Aβ_1-40_, pyroglutamated AβpE_3-42,_ and AβpE_3-40_) in the following ways: (1) Aβ42 has a higher content of regular secondary structure (β in solution and α + β in membranes), (2) it is the only variant for which Tyr10 side-chains are inaccessible in either environment, (3) Aβ42 decreases the mobility of membrane lipids whereas the other three increase lipid mobility, (4) Aβ42 forms more regular discrete single channel conductances in lipid bilayers, and (5) atomic-force microscopy images of membrane-embedded Aβ42 assemblies are more channel-like ((Karkisaval et al., 2024)).

The hydrophobic alkyl phase of membranes likely increases the secondary structure of the peptides’ transmembrane regions. But sizes of Aβ42 assemblies vary enormously in both aqueous and membrane environments and their shapes change gradually with time. Nonetheless, we have modeled this range of assemblies with a common structural motif, i.e., the last third of the peptide (S3) always forms a hydrophobic antiparallel β-barrel that is either buried in soluble assemblies or exposed to lipid alkyl chains in the membrane. In solution this barrel is typically surrounded by an outer β-barrel formed by the more hydrophilic first two-thirds of the peptide (segments S1 and S2). For channel models of this paper, all Aβ42 subunits have identical conformations, S2 segments form amphipathic α-helices on the membrane surfaces and S1 segments form two in tandem, pore-lining β-barrels inside the S3 β-barrel. Models of the accompanying manuscript have two unique peptide conformations; half of the S2 segments form part of a pore-lining β-barrel and the other half form surface α-helices.

Analyses of Aβ sequence homology suggest that some Aβ assemblies are highly structured and functionally important. For example, amyloid β sequences of humans, snapping turtles, and finches are identical and a sequence from the bamboo shark has only one difference, a conservative I31V mutation. Greater differences occur in bony fish, but conservation of S3 is still remarkable (Fig. 1). This high degree of conservation is consistent with findings that Aβ assemblies are functionally important ((Fagiani et al., 2021), (Brothers et al., 2018), (Bartley et al., 2022), (Pannuzzo, 2022)) and have highly ordered structures or family of structures under some conditions. Functional structures are unlikely to be fibrils, which are typically considered to be deleterious.

**Figure 1.**
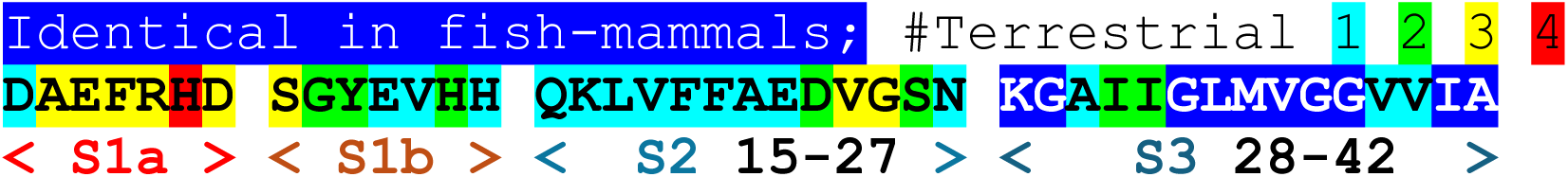
Results of a multisequence alignment for Aβ42. Positions identical among all vertebrate sequences are dark blue, the remainder colors indicate the degree of conservation among terrestrial vertebrates (amphibian - mammals): cyan identical in all, green 2 residue types, yellow 3 types, red 4 types. Our nomenclature divides the sequence into three segments of equal length: S1a is the least conserved, three of the last four positions in S1b are conserved, positions Q15 - E22 in S2 are conserved, and ten of fifteen positions in S3 are identical among all species including fish. The sequence variability correlates with the segment polarities; S1a is the most hydrophilic and S3 is the most hydrophobic.

Our models are hypothetical because current experimental data are insufficient to solve the structures directly. Some results, however, support aspects of our models. For example, S2 segments form antiparallel β-strand pairs in some of our models of soluble oligomers. This also occurs in at least one fibril structure for the Iowa mutant (Fig. 2A) ((Qiang et al., 2012)). In some of our models S2 and S3 have a U-turn structure resembling those of the antiparallel and some parallel fibrils (Fig. 2B). An A2T mutation is protective whereas A2V, H6R, and D7H, ((Aggarwal & Biswas, 2021)) are pathogenic. GM1 does not bind to nontoxic rat Aβ which has mutations at S1 positions R5, Y10, and H13 ((Foroutanpay et al., 2018)). Also, the structure of AβpE_3-42_ differs from that of Aβ42 (Karkisaval et al., 2024). Additionally, these findings suggest that S1a is important for the function of Aβ42 assemblies and should be included in structural models. Unfortunately, S1 is the most difficult segment to model; it is disordered in numerous fibril and NMR-based structures. However, in some structures it has a β secondary structure. Fig. 2 C-E show that S1 can form an extended β-strand ((Soldner et al., 2017; Staff, 2017)), two perpendicular strands ((Kollmer et al., 2019)), or a β-hairpin ((Bakrania et al., 2022)). Similar structures are present in our models.

**Figure 2.**
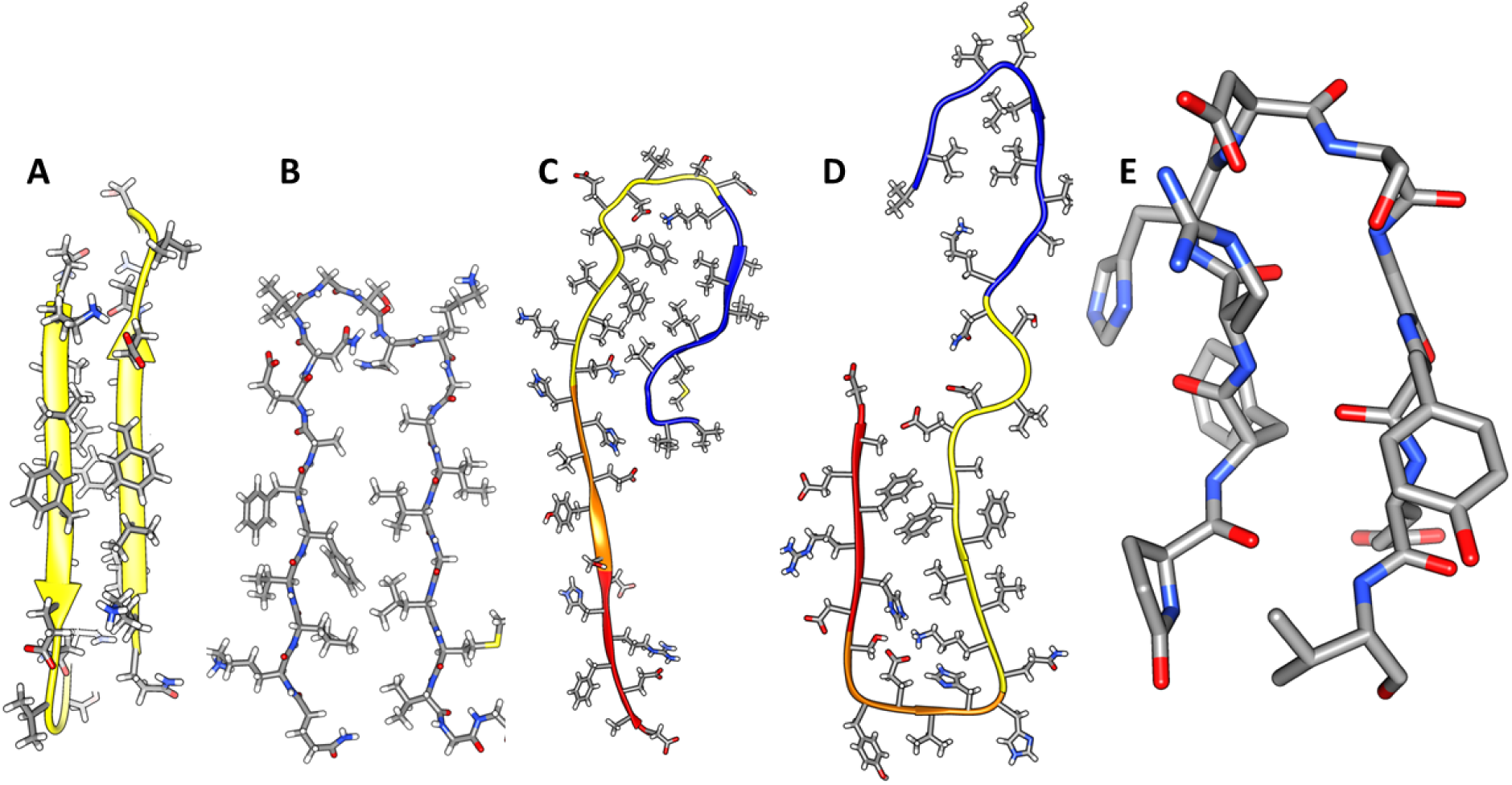
Experimentally-determined Aβ structures with features incorporated in our models. Side-chains are colored by element (carbon – grey, nitrogen - blue, oxygen – red, sulfur – yellow, hydrogen – white). (A & B) Front views of exposed faces of antiparallel S2 strands and U-turn β structure of S2 (left) and S3 (right) segments of an Iowa mutant fibril ((Qiang et al., 2012), PDB ID: 2LNQ). (C) Portion of a monomer structure from a fibril in which S1 forms a continuous β-strand ((Soldner et al., 2017), PDB ID: 2M4J). (D) Another monomer structure from a fibril in which S1a and S1b form different β-strands ((Kollmer et al., 2019), PDB ID: 6SHS). Hydrophobic side-chains of S1a (red backbone) pack next to hydrophobic side-chains on the more polar face of S2 strands (yellow backbone). S1b strands (orange backbone) connect S1a to S2. (E) Structure of a S1a-S2b β-hairpin bound to an antibody ((Bakrania et al., 2022), PDB ID: 7OW1 & 7OXN).

The following additional results support our models: (1) The structure of the 6-stranded antiparallel S3 β-barrel at the core of our hexamer models resemble that of Cylindrin ((Do et al., 2016)). (2) A β-barrel shaped Aβ42 structure forms in detergents ((Osterlund et al., 2019)). (3) NMR studies of Aβ42 channels indicate that they form well-ordered β-barrels with no more than two subunit conformations ((Serra-Batiste et al., 2016)). (4) Aβ oligomers have antiparallel β secondary structure ((Cerf et al., 2009), (Vignaud et al., 2013)). (5) Our models were constructed to be consistent with results of atomic-force microscopy ((Quist et al., 2005), (Connelly et al., 2012)) and freeze fracture microscopy images ((Shafrir, Durell, Arispe, et al., 2010)). Additional supporting findings for specific models are presented below.

## METHODS

Most methods used to develop our models were the same as those reported previously ((Durell et al., 2022)). Briefly, sequences of Aβ homologs were collected and aligned using the Blastp program (https://blast.ncbi.nlm.nih.gov > Blast.cgi). For 3D structural models, β-barrel parameters were calculated using theory developed by Murzin *et al*. (Murzin et al., 1994) and Hayward and Milner-White (Hayward & Milner-White, 2017). Gap distances between the walls of the adjacent barrels were constrained to be between 0.6 and 1.0 nm in the absence of lipids and between 1.0 and 1.4 nm if GM1 alkyl chains bind between the barrels. We selected models that maximize hydrogen bonds, salt-bridges, interactions among aromatic side-chains, burial and tight packing of hydrophobic side-chains that are highly conserved among Aβ homologs, and aqueous solvent exposure of hydrophilic side-chains, especially for positions that are hypervariable among homologs. Residues with polar side-chains and/or that have a high propensity for turn or coil secondary structure (Chou & Fasman, 1978) were favored for connecting loops. The final constraints were experimental: the sizes, shapes, molecular weights, secondary structures, and EM images of Aβ42 assemblies. All atomic-scale structures were generated “manually” with the UCSF Chimera program (Pettersen et al., 2004), with the aid of an in-house program to create the β-barrel structures. These were iteratively subjected to energy-minimization using the CHARMM program (Hwang et al., 2024) to eliminate atomic overlap, and optimize covalent geometry and salt-bridges and hydrogen bonds. Once these initial models had been developed, each was subjected to 200 ns of molecular dynamics (MD) simulation.

Some models were subsequently adjusted, remade symmetric and energy-minimized, and then resubjected to MD simulations. The final structures were deemed stable when all-atom Root-Mean-Squared-Deviations (RMSDs) over the trajectories were below a threshold of 5.0 Ẩ. In some trial models, portions of soluble domains and of GM1 gangliosides never stabilized; most of these were rejected.

The MD simulation input files were prepared using the Charmm-GUI web-based service (Jo et al., 2008; Lee et al., 2016; Lee et al., 2019; Park et al., 2021). Water soluble models were processed using the Solution Builder while transmembrane models were processed using Membrane Builder. The membrane was composed of 1-palmitoyl-2-oleoyl-sn-glycero-3-phosphoethanolamine (POPE) lipids. Gromacs 2024 (Bekker et al., 1993) was used to run the simulation on the NIH High Performance Biowulf cluster (https://hpc.nih.gov).

## RESULTS

Our simplest β-barrel models of Aβ42 are composed of N identical subunits arranged with N/2-fold radial symmetry about the barrels’ central axes and P2 symmetry (180^◦^ rotation about axes perpendicular to and that intersect the radial axis at the assembly’s center of mass).

### Soluble oligomers

Our mission is to not only understand structures of some Aβ42 assemblies but also to understand processes by which they assemble in the aqueous phase, interact with membrane surfaces, penetrate membranes, and form ion channels. Although we propose detailed models for some soluble assemblies, we realize that these depictions are likely overly-structured and may be relatively unstable. In fact, if they were too stable the kinds of transitions and morphing we propose would be improbable.

In our analysis of Aβ42 beaded annular protofibrils (bAPFs), we identified five sizes of beads which we modeled as hexamers, octamers, dodecamers, hexadecamers and octadecamers (Fig. 3). Although many bAPFs have irregular shapes and are composed of multiple sizes of beads, we identified 29 subsets of symmetric images of bAPFs; each image was composed of the same size and number of beads. Averaged images of subsets composed of putative hexamers and dodecamers are shown in Fig. 3. Circular APF assemblies range in diameter from ∼4 to 50 nm.

**Figure 3.**
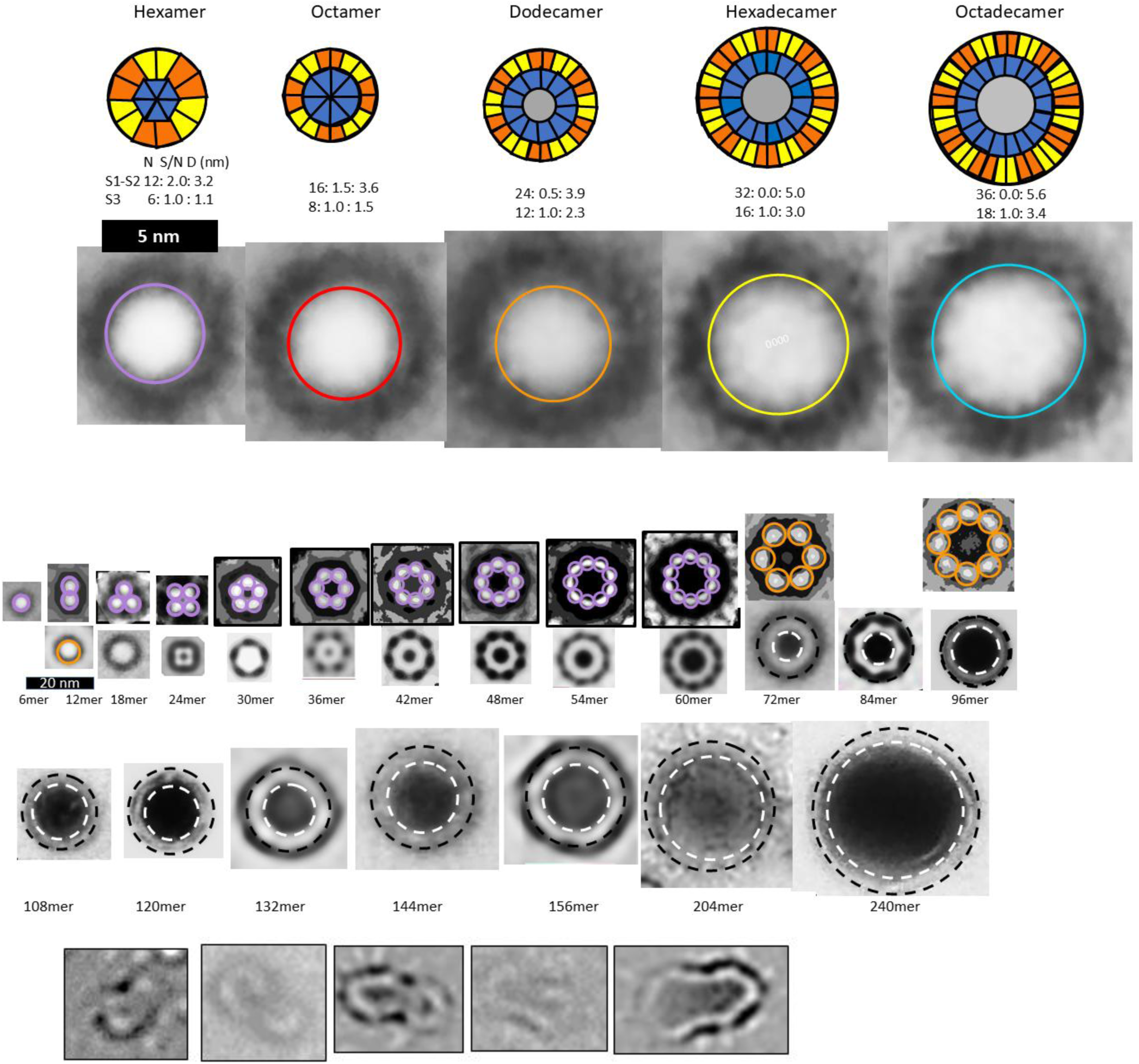
Averaged images of beaded and smooth APFs (adapted from (Durell et al., 2022)). Top row: Wedge schematic models of five soluble Aβ42 oligomers showing relative positions of segments in a middle cross-section of the β-barrels; inner S3 β-barrels are blue, outer S1-S2 β-barrels are orange and yellow, gray center circles represent hexane or lipids inside larger S3 barrels. Number of subunits (N), sheer number/N ratio (S/N), and diameter of barrel backbone (D) parameters of each β-barrel are below schematics. Second row: APF images of isolated beads. Circles have the same diameter as the outer S1-S2 barrels of the schematics. Third row: assemblies of putative hexamer (purple) and dodecamer (orange) beaded APF. The next two rows are images of smooth APFs.

Approximated numbers of Aβ42 peptides in each assembly are indicated below the images. The apparent wall thickness is about the same for all the larger sAPFs as indicated by the dashed lines. The bottom row has images of APFs that may be merging or dividing.

Fig. 3 is shown to make the following points: (1) Soluble oligomers proposed to comprise bAPFs can have multiple sizes. (2) Clusters of soluble beads in bAPFs, each with its own set of concentric β-barrels in our models, gradually morph into smooth annular protofibrils (sAPFs), which have only one set of concentric β-barrels. (3) Diameters and the number of monomers comprising sAPFs vary greatly, but the apparent thickness of their walls remains relatively constant. (4) We propose that Aβ42 assemblies can transition from one size to another; *e.g.*, two hexamers may merge to form a dodecamer, two dodecamers may merge to form a 24mer, two 24mers may merge to form a 48mer, *etc*., and (5) we propose here that similar processes occur for transmembrane Aβ42 assemblies.

### Soluble hexamers

The most frequently observed bead size in our analysis of annular protofibrils corresponded to our model of soluble hexamers that was developed before our analysis of APFs. We have developed two models of soluble hexamers: a small hexamer composed of only Aβ42 subunits (Fig. 4) and a larger hexamer that includes six GM1 gangliosides (Fig. 5). Established β-barrel theory ((Murzin et al., 1994), (Hayward & Milner-White, 2017)) constrain the tilts of β-strands for symmetric models. If all antiparallel S3 strands have identical conformations then the antiparallel S3 barrel can have only two tilt angles, α = 36^◦^ if S/N = 1.0 for the small, 1Con1 models (Fig. 4) or 55^◦^ if S/N = 2.0 for the large, 1Con2 models (Fig.5) see Supplement Fig. S1.

**Figure 4.**
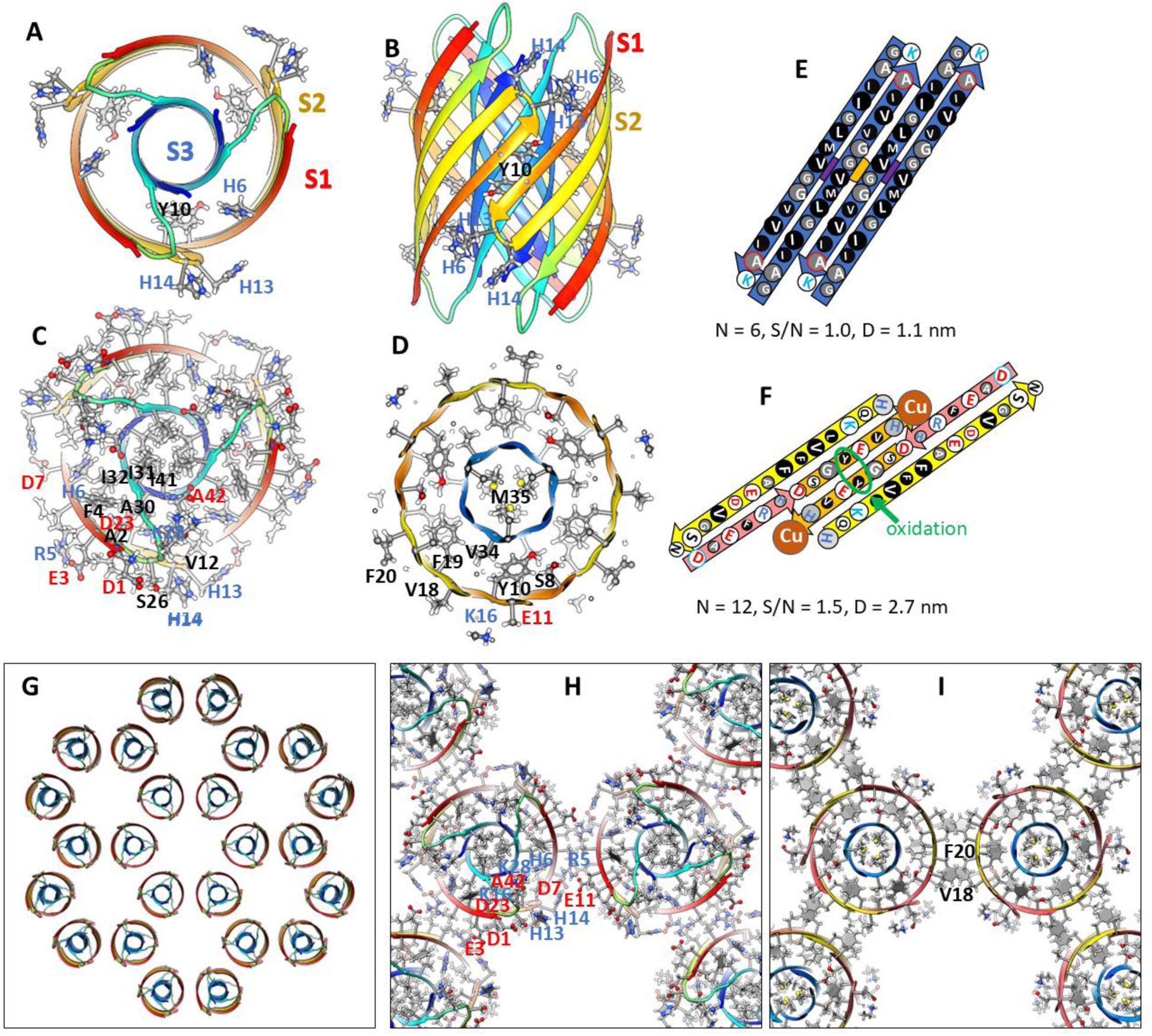
Model of a small 1Con1 soluble Aβ42 hexamer and hexagonal lattice of soluble hexamers. (A & B) Top and side views of backbones, histidine and Tyr10 side-chains are labeled. The backbone is represented by rainbow-colored ribbons (red at N-termini to blue at C-termini). (C & D) Cross-sections of the ends and mid-region of the models with side-chains. Oxygen, nitrogen, and sulfur atoms have larger diameters and are colored red, blue, and yellow; gray aromatic rings are filled. Residues of one subunit are labeled. (E &F) Flatten representation of four subunits for the inner S3 and outer S1-S2 β-barrels showing relative positions of the side-chains. Barrel parameters are below schematics. Arrows represent proposed β-strands, red for S1a, orange for S1b, yellow for S2, and blue for S3. Larger circles indicate side-chains on the exterior of the barrels, smaller circles are side-chains on the interior. Side-chains are colored by their polarity: white-blue letter = positively charged, white-red letter = negatively charged, white-black letter = uncharged polar, gray-blue letter = Histidine, gray-black letter = ambivalent, black-white letter = hydrophobic. Purple and orange circles behind the S3 schematic indicate axes of P2 symmetry. (G-I) Hexagonal lattice of 24 soluble hexamers. (G) Backbones. (H & I) Enlarged portion showing contacts between hexamers with side-chains for top and middle cross-sections.

**Figure 5.**
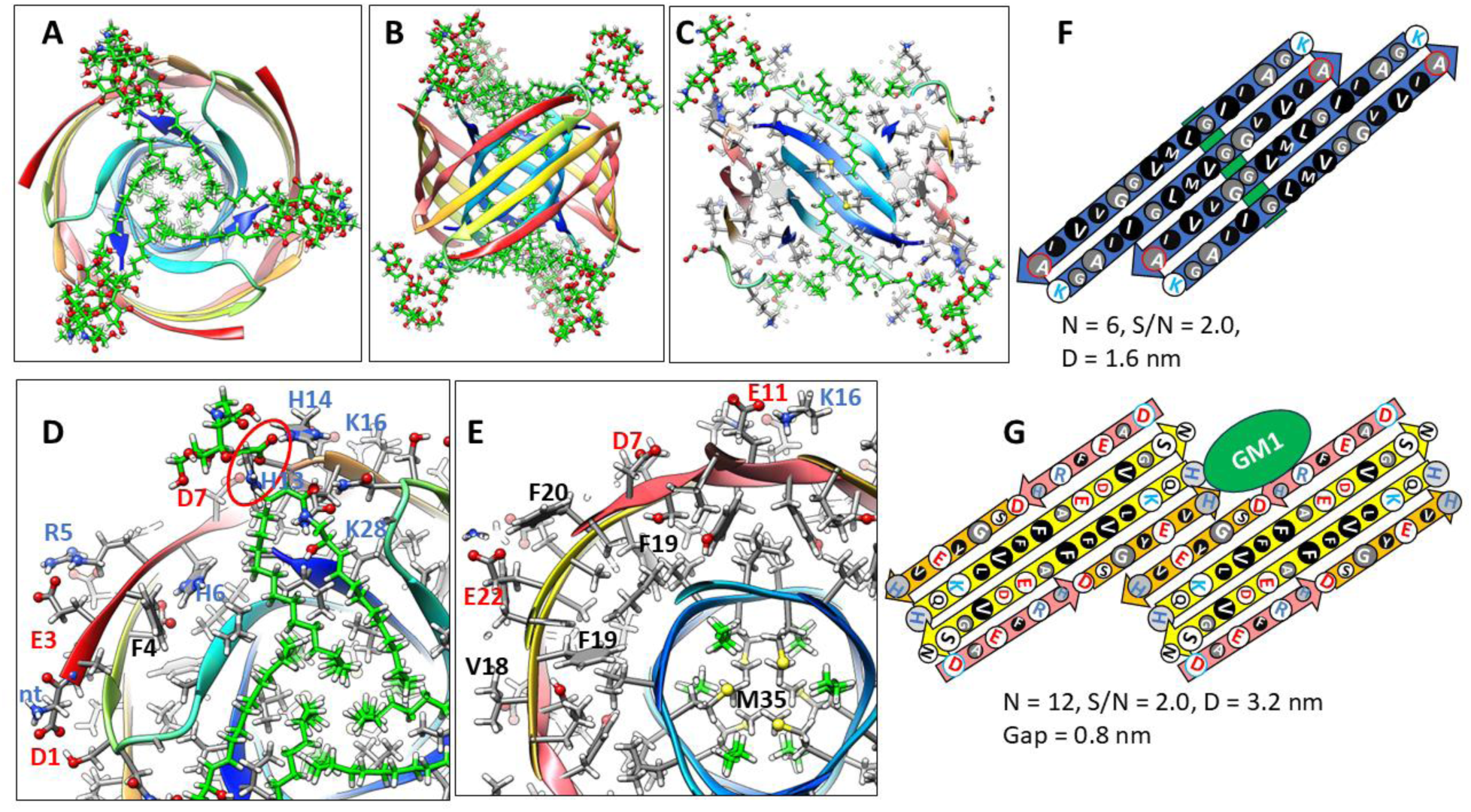
1Con2 model of a soluble hexamer with bound GM1 gangliosides. GM1 carbons are green, other atoms are colored by element. Aβ42 segments are colored as in Fig. 4. (A & B) Top and side-views of the backbone rainbow-colored ribbon and GM1 gangliosides with alkyl chains inside the S3 β-barrel. (C) Cross-section of the S3 barrel from the inside showing GM1 alkyl chain (green) packing next to inwardly-facing Gly33 and Gly37 of the Glycine pleat. (D & E) Portions of (D) top and (E) middle radial cross-section with side-chains and GM1 gangliosides. The GM1 carboxyl group is encircled in red and binds to side-chains of His13 and His14. Other side-chains are labeled for one subunit. (F & G) Flattened schematics of inner S3 and outer S1-S2 β-barrels with parameters. (F) A Glycine pleat is highlighted with green background and (G) location of GM1 headgroup is indicated by a green oval.

Previously we developed and analyzed multiple concentric β-barrel models of the small soluble Aβ42 hexamer; we favored models in which all subunits of antiparallel models have identical structures and interactions and are related by 3-fold radial and 2-fold perpendicular (P2) symmetry ((Shafrir, Durell, Anishkin, et al., 2010),(Yun et al., 2011)). Two NMR structures of Aβ42 oligomers, a tetramer ((Ciudad et al., 2020)) and a 32mer ((Gao et al., 2020)), conclude that pairs of S3 β-strands are antiparallel with a P2 axis of symmetry between the Val36 residues. This feature exists in our current small hexamer model (Fig. 4) and was present in our 2010 model (Shafrir, Durell, Anishkin, et al., 2010). The model of Fig. 4 resembles our previous models: six antiparallel S3 strands comprise a core antiparallel β-barrel that has no polar side-chain atoms, odd-numbered side-chains are in the interior. A surrounding β-barrel formed from S1 and S2 β-strands shields the S3 core from water. The position of S1 relative to S2 was altered in the current model to make it more consistent with new data on the formation of a Cu^2+^ ((Williams et al., 2016)) binding site and cross-linking of monomers by oxidation of the Tyr10 side-chains ((Maina et al., 2023), (Urbanc, 2021)) (Fig. 4F).

Six S1-S2 β-hairpins comprise a second β-barrel that surrounds the S3 barrel in these models. The models of Fig. 4 (S/N = 1.5) and 5 (S/N =2.0) were used for these models because they predict a near optimal gap distance of ∼0.8 nm between the walls of the outer and inner barrels in both models.

Most hydrophobic side-chains of these models are buried except for Val18 and Phe20 in the center of the S2 strand. These hexamers can aggregate to form a hexamer-of-hexamers (the most frequently observed bAPF assembly of Fig. 3) or a larger hexagonal lattice with regions of 2-fold, 3-fold, and 6-fold radial symmetry (Fig. 4 G-I). If this occurs, Val 18 and Phe20 of adjacent hexamers interact at the region of 2-fold radial symmetry and thus become buried in the lattice models (Fig. 4I).

Changing the tilt angle of the S3 strands of the 1Con2 of Fig. 5 to 55^⸰^ increases the diameter of the S3 barrel and allows Gly33 and Gly37 residues to align on the same pleat. The absence of side-chains in these pleats creates space for one alkyl chain of each GM1 lipid to pack next to the wall inside the S3 barrel (Fig. 5 E & F). The GM1 headgroups extend into the aqueous phases where their carboxyl group binds to the His13 and His14 side-chains. Tilts of S1 and S2 strands are also increased to 55^⸰^ to make the S1-S2 barrel large enough to surround the S3 barrel.

### Soluble dodecamer

S3 barrel diameters increase for larger soluble oligomers. This raises the possibility of their hydrophobic interiors being filled with hydrophobic molecules (e.g., lipids, fatty acids, detergents, hexane). In the soluble dodecamer model of Fig. 6, alkyl chains of GM1 lipids fill the inside of the 12-stranded antiparallel S3 β-barrel. S1-S2 strands form an exterior 24-stranded antiparallel β-barrel that shields the S3 barrel’s outer surface from water. GM1 head groups extend into the aqueous phase and interact with the His13-His4 region linking S1 to S2.

**Figure 6.**
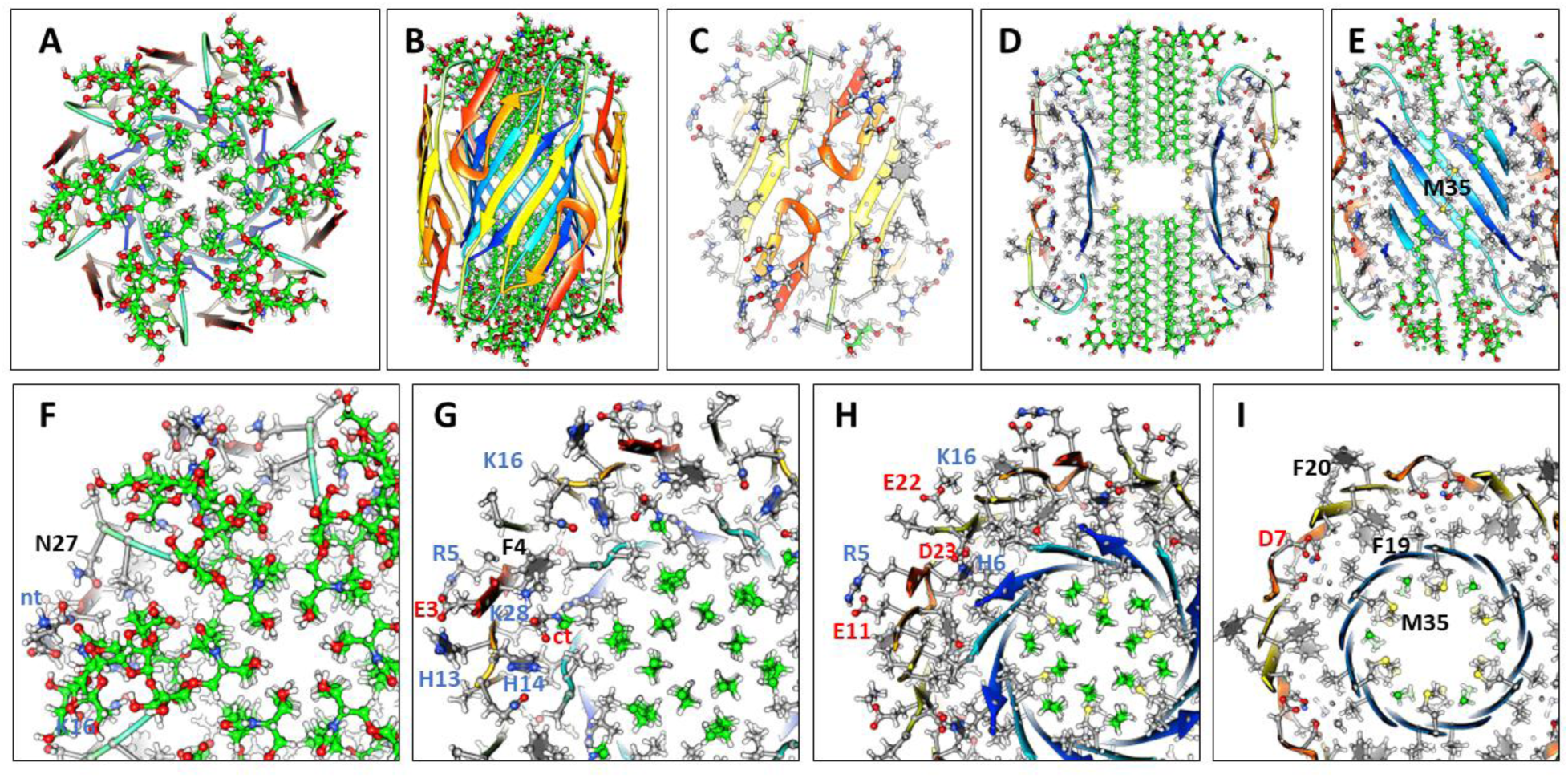
1con1 model of a soluble dodecamer with bound GM1. (A & B) Top and side views of the backbone and GM1 gangliosides. (C) Side view of outer S1hairpin (hp)-S2 barrel with side-chains. (D) Side cross-section showing GM1 alkyl chains inside the S3 barrel. (E) Side cross-section showing how alkyl chains pack next to the S3 barrel. (F-I) Portions of serial radial cross-sections beginning at the top and ending in the middle; some side-chains of one subunit are labeled.

### Models of Transmembrane Oligomers (TMOs) and Channels

#### 1Con models with a multiple of six subunits

Our working hypothesis is that soluble oligomers bind to membrane surfaces, then cluster before penetrating the membrane to form transmembrane oligomers (TMOs). Dodecamer and octadecamer TMOs may form either from the merger of two or three hexamer TMOs or from insertion of soluble dodecamers and octadecamers into the membrane. Clusters of TMOs may morph into channels that have only one S3 barrel (Fig. 7) in a manner analogous to how bAPFs morph into smooth APFs (Fig. 3). Sizes and shapes of these models are consistent with freeze-fracture images of transmembrane Aβ42 assemblies. These models have the following features in common: They all have an integer multiple of six subunits and all subunits within an assembly have the same conformation and interactions. S3 segments form membrane-spanning antiparallel β-barrels that interact with lipid alkyl chains. S1-S2 segments of TMOs extend into aqueous phases on each side of the membrane. S2 segments of channels form amphipathic α-helices on membrane surfaces; whereas, S1 segments form short β-barrels on each side of the membrane that stack end-on to line pores inside the S3 barrels. Freeze-fracture images with apparent pores in their centers have sizes consistent with channels possessing 12 to 36 subunits. Diameters, D, of putative 1Con β-barrels depend both on the number of subunits, N, and the tilt angle, α, of the strands; for a given N, D is smaller if S/N = 1 (α = 36⸰ and D = 0.19N) than if S/N = 2 (α = 36⸰ and D = 0.27N) (see supplement Fig. S1).

**Figure 7.**
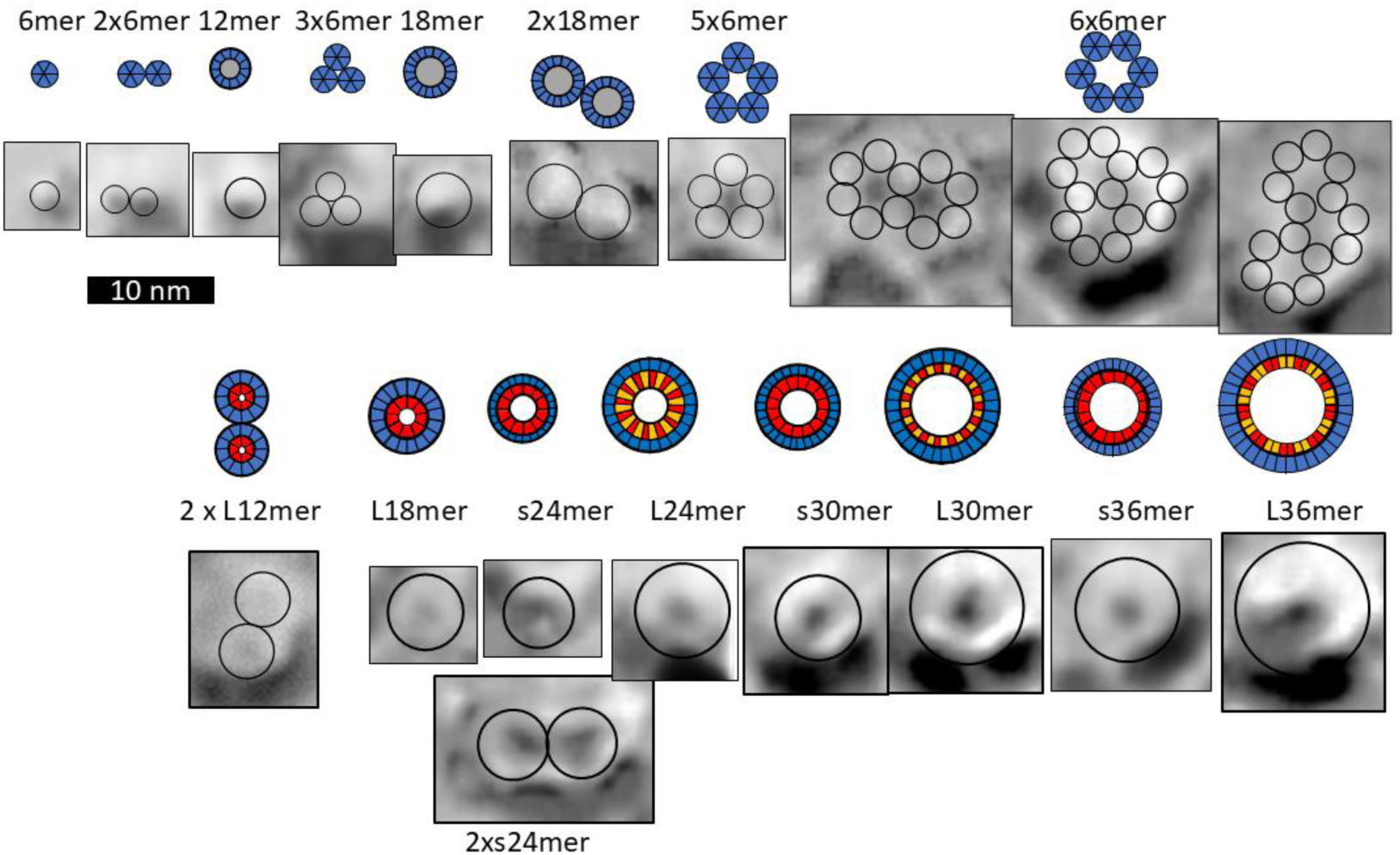
Schematics of β-barrel models composed of a multiple of six Aβ42 subunits plus freeze-fracture (FF) images (Shafrir, Durell, Arispe, et al., 2010) of putative Aβ42 transmembrane oligomers and channels. The top row shows schematics of S3 β-barrels for TMO assemblies for hexamers, dodecamers, and octadecamers. FF images with sizes and shapes consistent with the schematics are in the second row. Circles in the FF images have the same outer diameter as those of the schematics. Assemblies with five or six hexamers may have a central pore. Hexamer-of-hexamers may overlap to form partial hexagonal lattices with two or three pores (last three images). The next rows show schematics and FF images for putative transmembrane channels. Red regions inside the blue S3 barrels represent polar pore linings comprised of parallel S1a β-barrels. Larger barrels with the same number of subunits have linings composed of S1a (red)-S1b (orange) β-hairpins. S3 strands of the smaller barrels are less tilted (S/N = 1.0) than those of the larger barrels (S/N = 2.0).

Numerous FF images larger than those of Fig. 7 have no apparent pore through their center (Fig. 8). We propose that these have morphed from larger clusters of 12mers and/or 18mers. Our models of these larger assemblies have two S3 β-barrels; subunits on the exterior of the clusters (outside the red lines in schematics of Fig. 8) comprise an outer S3 barrel that surrounds an associated S1 barrel and those on the interior of the clusters comprise an inner S3 lipid-filled barrel that is surrounded by an S1-S2 barrel, as proposed for soluble assemblies. We call these plugged channels; however, in some cases the inner barrels may be extruded into the aqueous phase leaving relatively large channels with 32 to 72 subunits while creating soluble Aβ-lipoproteins observed in the aqueous phase (Michikawa et al., 2001).

**Figure 8.**
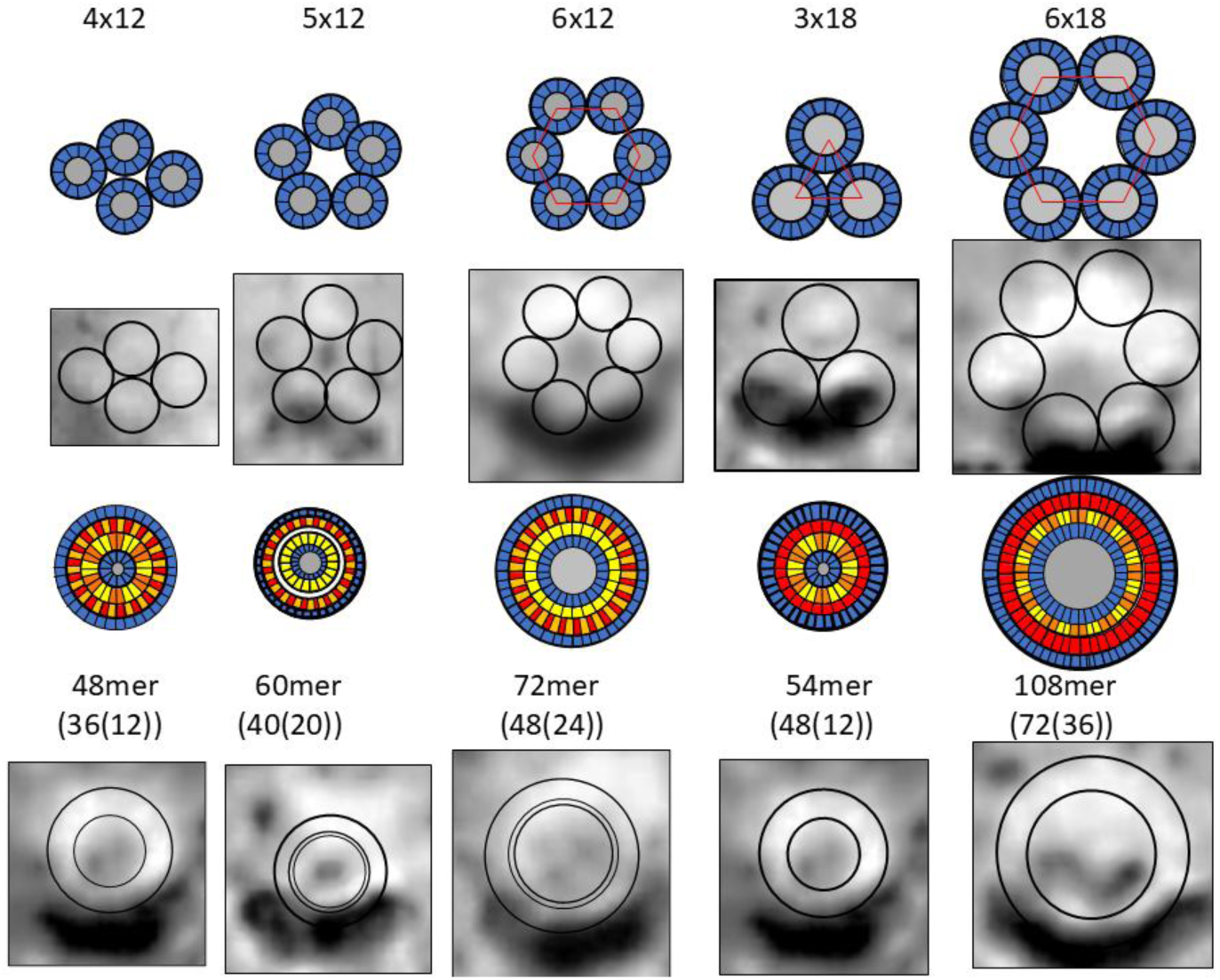
Schematics and FF images for putative plugged channels. (Rows 1 and 2) Assemblies of dodecamer and octadecamer TMOs. Gray regions in the centers of the schematics represent lipids. These assemblies may morph into the blocked pores in rows 3 and 4. Numbers of subunits in the outer channel-like and inner lipoprotein-like assemblies are in parentheses. Circles in the FF images have the outer diameters of the proposed channel S3 barrels and lipoprotein S1-S2 barrels.

In Aβ42-GM1 assemblies isolated from liposomes, S2 can be cleaved by trypsin whereas S1 and S3 cannot (Zhang et al., 2022). These assemblies range in molecular weight from 45–100 kDa*; i.e.*, are composed of ∼ 10 - 22 peptides. Based on these findings and the sequence of S2, we propose that S2 forms an amphipathic α-helix on the membrane surfaces when GM1 is bound to the channels as illustrated in the flattened schematics of Fig. 9.

**Figure 9.**
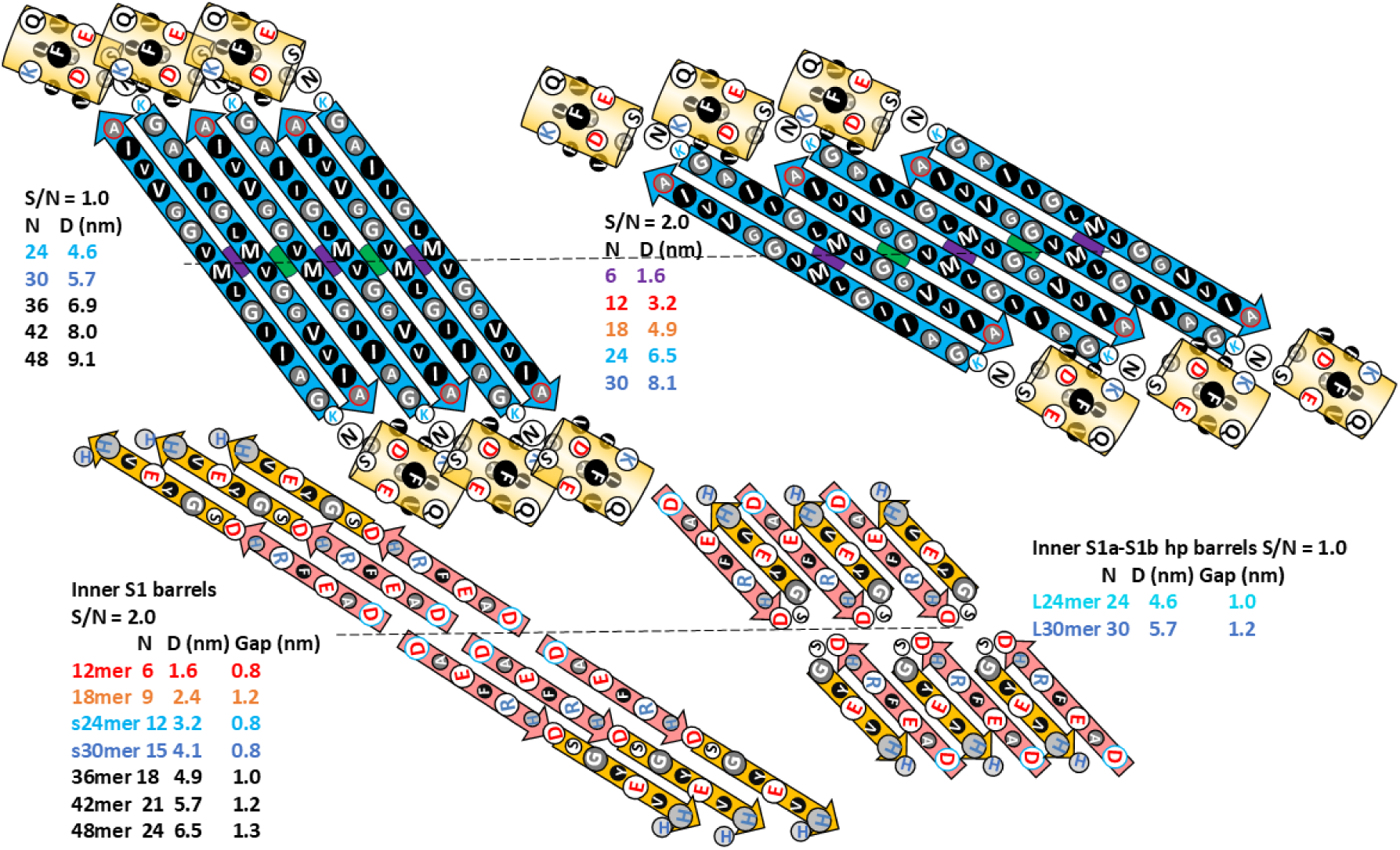
Flattened representation of six subunits for 1Con β-barrels viewed from inside the β-barrels. Dashed lines and purple and green circles indicate the plane and axes of P2 symmetry. Transmembrane assemblies of Fig. 7 can be formed by combinations of these motifs; parameters of each channel type are colored differently. Odd numbered S3 side-chains are oriented inwardly (larger circles). Most channels and outer blocked channels are lined by two S1 parallel β-barrels with S/N values of 2.0, but larger L24mer and L36mer channels are lined by two antiparallel S1a-S1b β-hairpin barrels.

### Membrane insertion mechanisms

We postulate that when soluble oligomers bind to a membrane’s outer leaflet, the hydrophobic S3 barrel remains intact, but some S1-S2 segments peel away and interact with lipid headgroups and the aqueous phase with their S2 segments becoming helical. Our funnel model of Fig. 10 is an aggregate of four hexamers. Most of the assembly has a cone shape that resides in the outer leaflet of the membrane; four S1 and two S2 β-strands form a β-sheet that shields the top portion of the tilted S3 barrel from water and two S2 α-helices bind on the sides of the S3 barrel. GM1 gangliosides bind between the hexamers where their headgroups bind or are near numerous histidine and arginine side-chains (Fig. 10C). The spout of the funnel penetrates the inner leaflet. It has a TIM-barrel type of α-β structure ((Wierenga, 2001)) in which eight S2 helices surround an 8-stranded parallel S1a β-barrel that lines a central pore.

**Figure 10.**
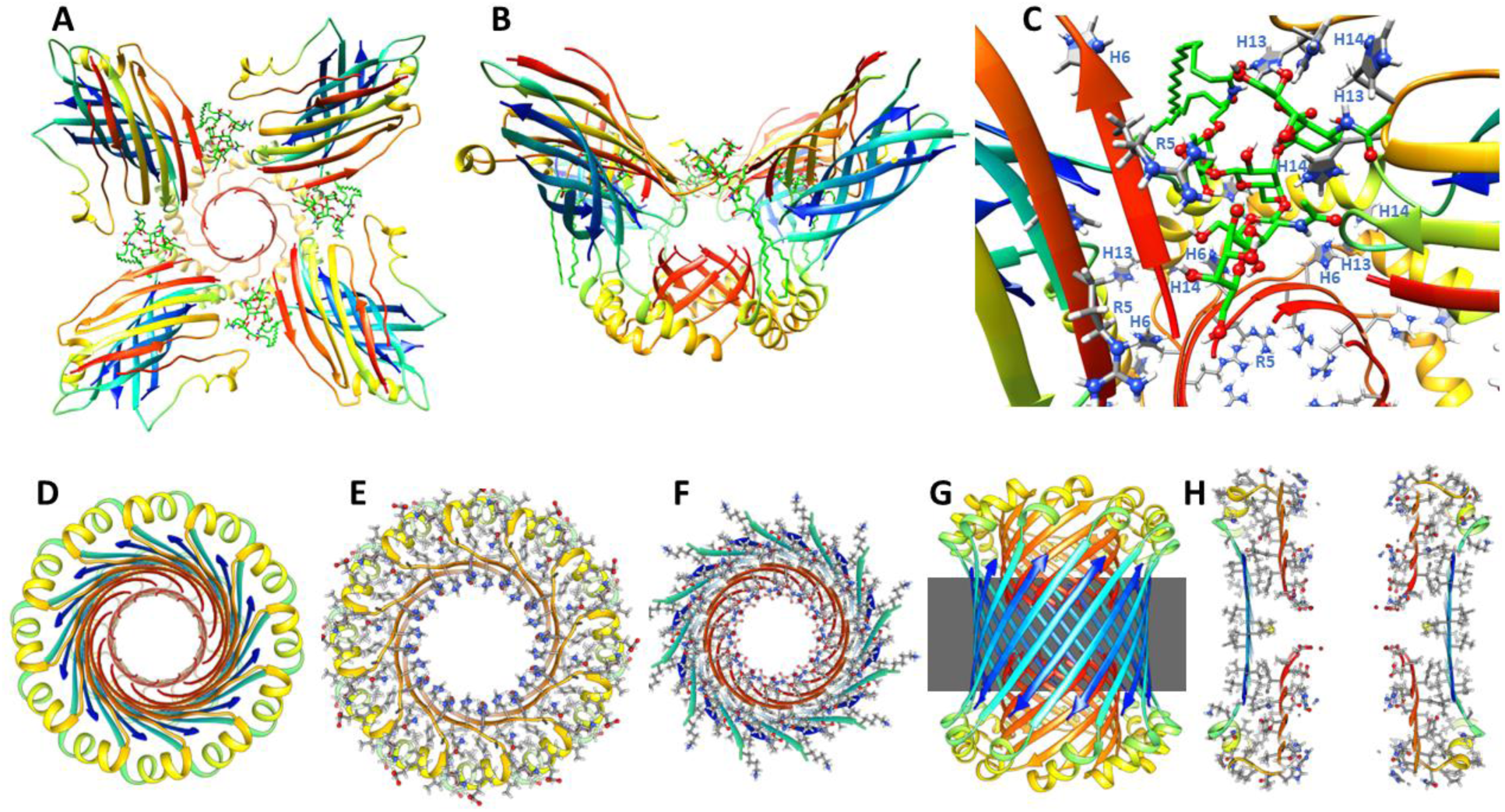
Atomic-scale models of the 4x6mer funnel model for aggregation and insertion of four hexamers through the membrane, and structure of a small 24mer channel proposed to develop from the hexamers. (A & B) Top and side views of four hexamers binding on the membrane surface with a TIM-like αβ-barrel penetrating the inner leaflet of the membrane and GM1 gangliosides binding between the hexamers. (C) Enlarged view of the GM1 head group in the putative binding site. Histidine and Arginine side-chains that interact with or are near the head group are labeled. (D-H) Small 24mer channel. (D) Top view of the 24mer channel backbone ribbon. (E & F) Cross-sections of soluble and transmembrane domains with side-chains. (G & H) Side views of backbone ribbon and cross-section with side-chains.

Funnel assemblies may morph into 1Con Aβ42 channels. We begin with models composed of 24mer subunits because they are consistent with recent experimental results obtained after our models were developed. Karkisaval et al, (2024) (Karkisaval et al., 2024) found that for membrane assemblies of Aβ42 the average orientation of β-strands with respect to the membrane normal was 30-40^⸰^, they had ∼46⁒ β-sheet and ∼19 ⁒ α-helix structure, and the α-helices were tilted 50-65^⸰^ relative to the membrane normal. In our small 24mer channel model S3 strands are tilted 36^⸰^, residues 2-13 and 31-41 are β (55⁒), residues 17-24 are helical (19⁒) and the helices are tilted by ∼55^⸰^.

Atomic force microscopy images of Aβ42 channels (Karkisaval et al., 2024) are fairly consistent with our models (Fig. 11) and appear more channel-like than assemblies of the other three variants. The diameter of the 24mer channel model’s soluble domain is ∼9 nm; the 4x6 funnel model is substantially larger. Most of the Aβ42 assemblies in the AFM studies extended from 1-3 nm above the membrane plane, the lower values are more consistent with our models.

**Figure 11.**
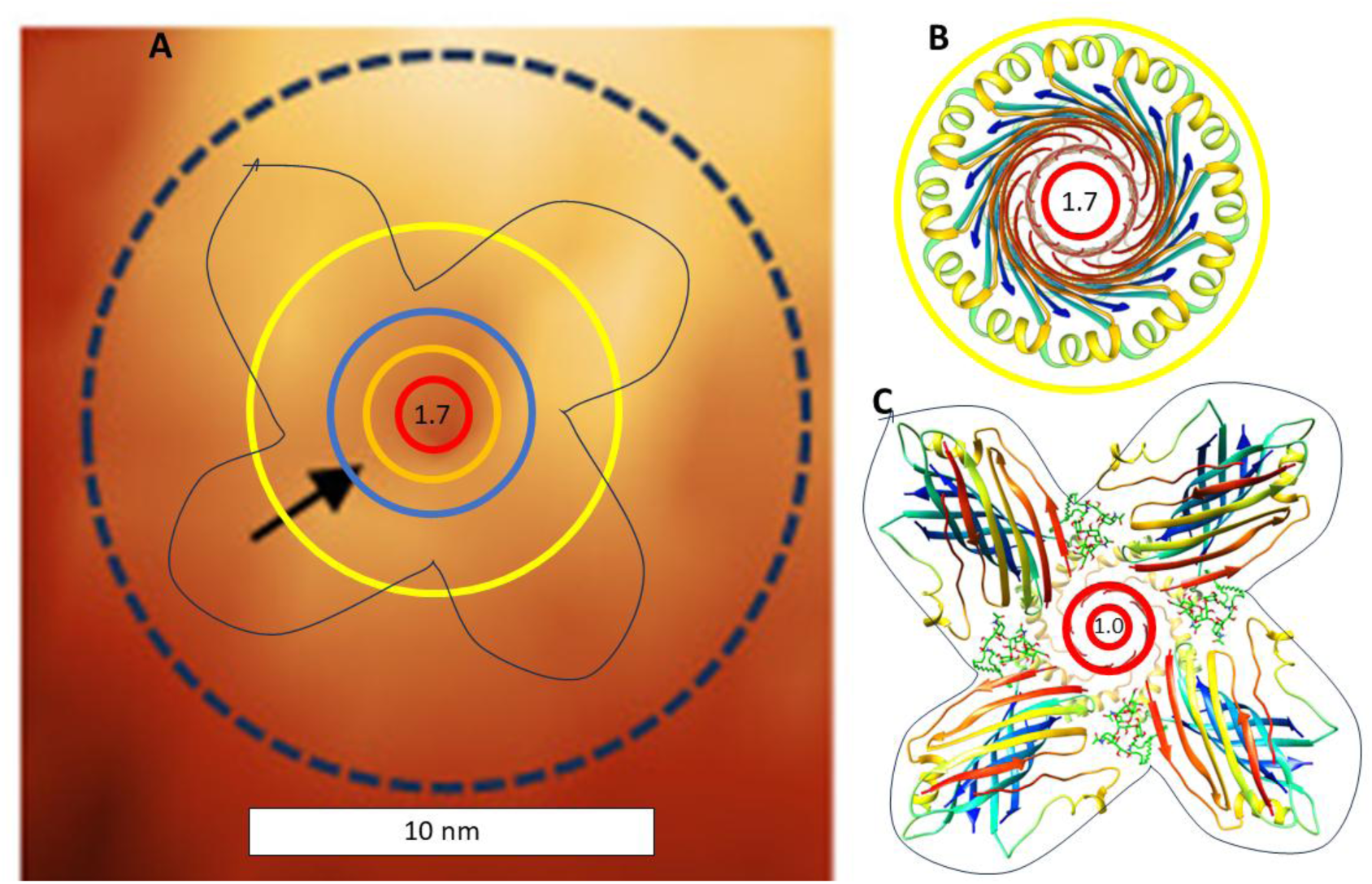
Comparison of the small 24mer channel model to recent results of atomic force microscopy of Aβ42 channels. (A) AFM image of Aβ42 channel with shapes and dimensions of the 24mer channel superimposed (adapted from Fig. 5i of (Karkisaval et al., 2024) http://creativecommons.org/licenses/by/4.0/). The added yellow circle is the outer edge of the small 24mer channel soluble domain, the blue circle represents the S3 backbone, orange circle represents the S1 backbone, and the red circle the size of the pore. The black outline represents the outer boundary for the 4x6mer-funnel insertion model. (B & C) Top view of the 24mer channel and insertion models with the same scale as the AFM image.

Based on the single channel conductance, Karkisaval et al., (2024) (Karkisaval et al., 2024) estimated the diameter of the Aβ42 pore to be 0.56 nm, about half and a third of those predicted by the inserting model and the channel model. In contrast, Bode et al. (2017) (Bode et al., 2017) estimated that Aβ42 channel pores in neurons have diameters of 1.7-, 2.1-, and 2.4-nm, more consistent with our models. Membranes of the Karkisaval et al, (2024) (Karkisaval et al., 2024) study contained no GM1 gangliosides. Fig.10 was developed for small hexamer and 24mer models. Similar models can be developed for large hexamers and 24mers (Fig. 12) proposed to bind GM1 gangliosides. Their presence in neurons may account for some differences observed in single channel conductances. S3 strands of the shorter-wider 1Con2 24mer model are more tilted (S/N = 2), the S2 helices are farther apart, the S1a-S1b β-hairpins form 24-stranded antiparallel barrels that line the larger pore, and the gap distance between the S3 and S1 barrels is greater. GM1 headgroups are modeled to bind between the S2 helices where their carboxyl group can interact with His 13 and His 14 (Fig. 12E) and their alkyl chains fit into the Glycine pleat between the S3 and S1 hairpin barrels in this model (Fig. 12H).

**Figure 12.**
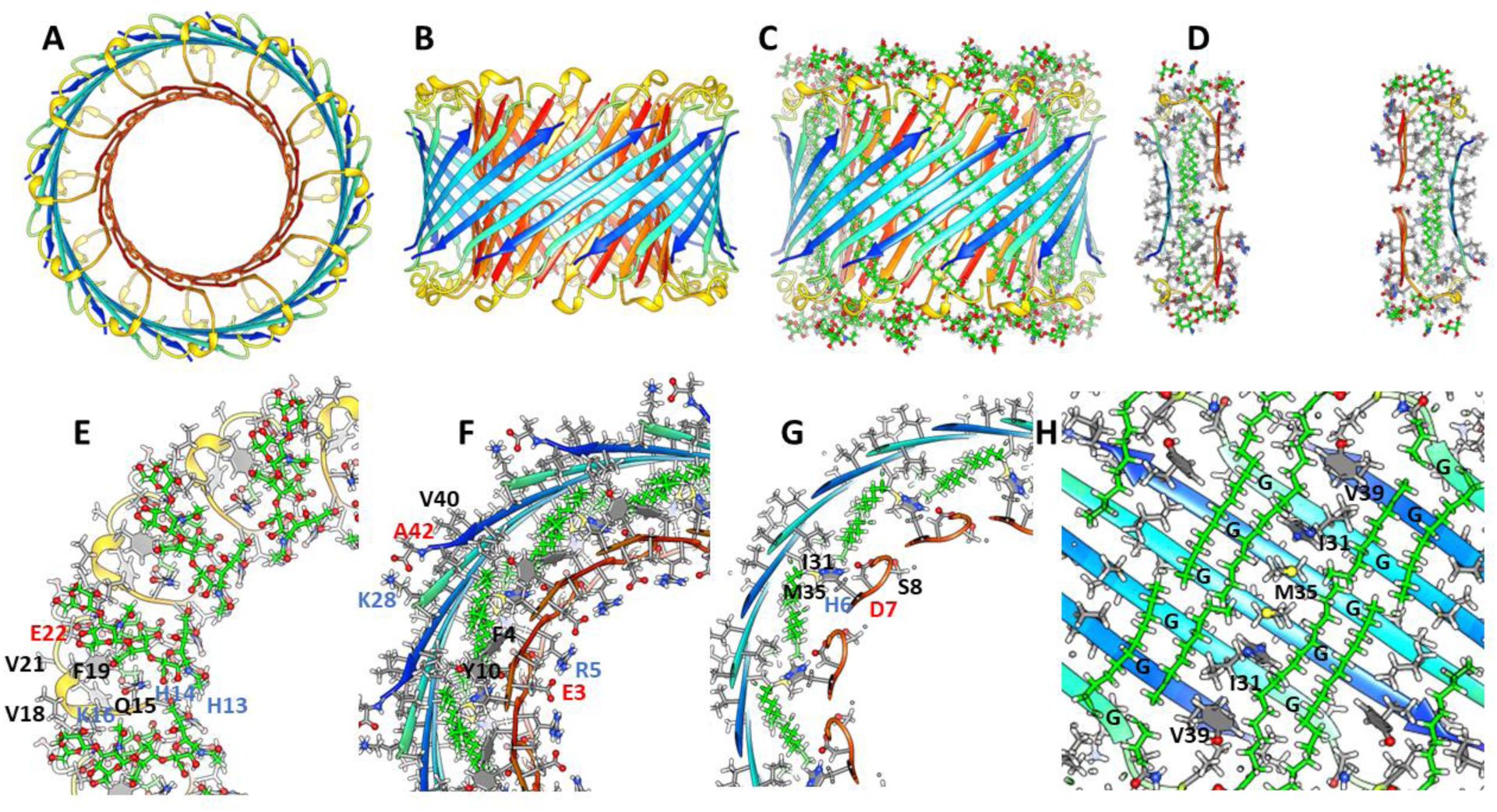
Atomic scale model of a large Aβ42 1Con2 24mer channel with GM1 gangliosides bound between the outer S3 and inner S1 β-barrels. (A - C) Top and side views of the Aβ42 backbone without and with GM1s (green carbons). (D) Clipped side view cross-section showing how GM1 alkyl chains fit between the outer and inner barrels and charged S1-hairpin side-chains line the pore. (E - G) Expanded portions of upper, next lower, and middle radial cross-sections with side-chains colored by element and important side-chains of one subunit labeled. (H) Clipped side view showing how GM1 alkyl chains fit in the Glycine pleats between I31, M35, and V39 side-chains.

### Models with six hexamers

Insertion of six hexamers into the membrane may involve formation of a 6x6mer funnel channel (Fig. 13 A&B). Next the S3 β-barrels may reorient to span the bilayer as an assembly of six hexamer TMOs. Atomic force microscopy studies of transmembrane amyloid assemblies indicate that the aqueous domains extend 2-5 nm above the surface of the membrane (Connelly et al., 2012). Assuming a standard membrane thickness of ∼ 5 nm, the soluble domains of the models should extend up 3-4 nm above the membrane’s surface. In the channel model of Figs. 13 E&F, S2 α-helices comprise a triple-stranded coiled-coil with side-chains of L17, V18, F20, and A21 comprising the hydrophobic core. Radially outward S1a-S1b β-hairpins of the 6x6 model interact in the aqueous phase to form a 24-stranded antiparallel β-barrel that surrounds the S2 coiled-coils. S1 segments of the subunits nearest the center interact to form 6-stranded parallel β-barrels with the S1a portion extending between the six S3 β-barrels from each side of the membrane to line a central pore with charged side-chains (Fig. 13 E&F). The packing among hydrophobic side-chains within the assembly is dense (Fig. 13H) and most charged side-chains are exposed to water and form salt-bridges. The soluble portion of this model is not well maintained during molecular dynamic simulation, but the transmembrane region is. Additional hexamers may become part of the assembly to form a hexagonal lattice (Fig. 13D) as supported by freeze-fracture images in Fig. 7.

**Figure 13.**
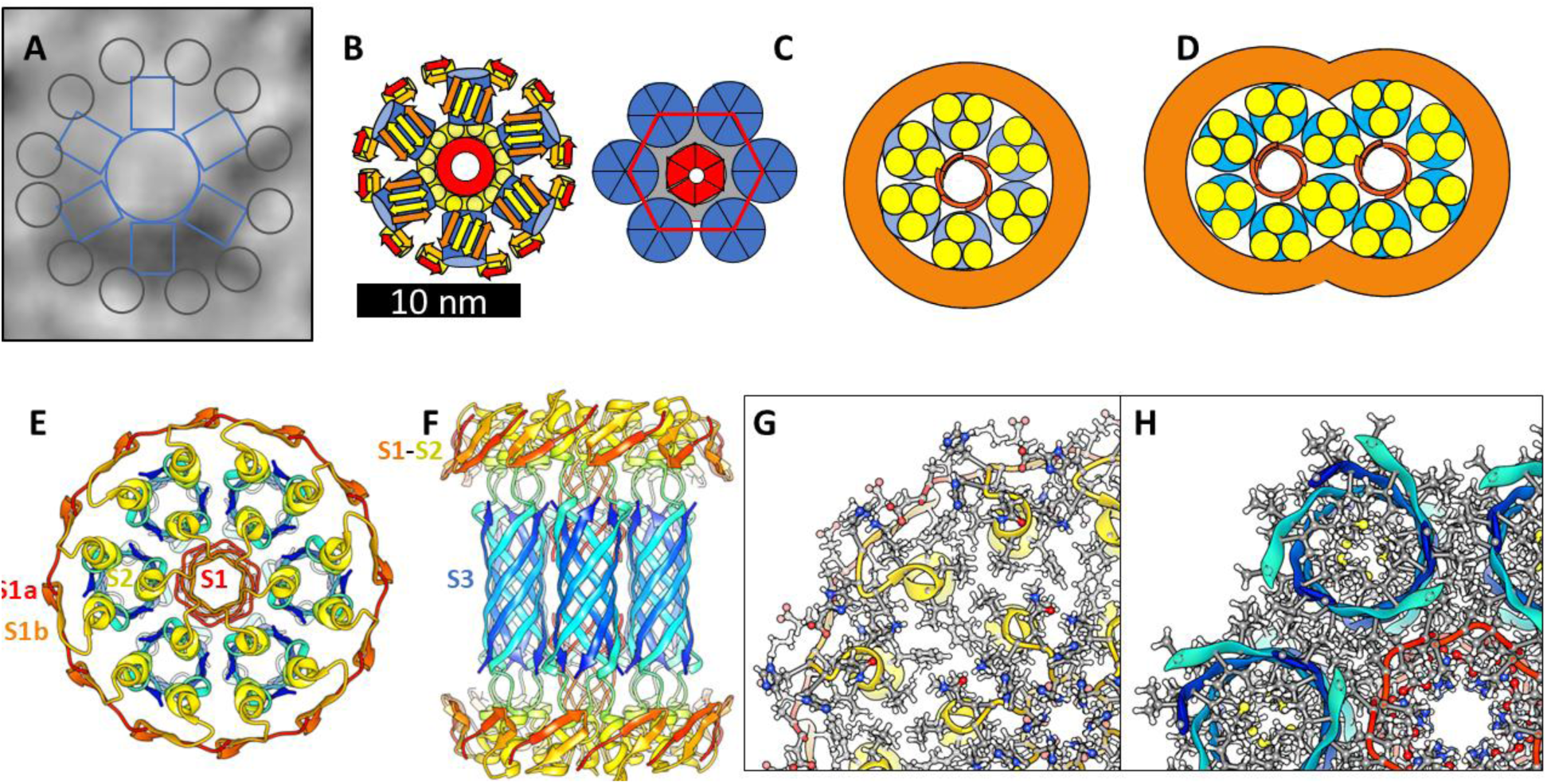
Models of hexamer-of-hexamers assemblies and freeze fracture image consistent with the surface models. (A) A large FF image with superimposed schematic outline of six radial arms each with two end globs extending from a central mass. (B) A schematic of six hexamers aggregated on the membrane surface. Blue S3 β-barrels comprise the bottom of radiating armes. A central region of S2 α-helices (yellow) surround a S1 β-barrel (red) that penetrates the inner leaflet of the membrane. The top portion of each S3 barrel (blue) is shielded from water by two antiparallel S1 (orange)-S2 (yellow) β-hairpins. S2 helices covered by S1 hairpins form an outer ring. (C) The hexamers reorient with S3 barrels spanning the membrane to form the 6X6mer channel. Aqueous phase S1-S2 segments are on the right. (D) A schematic of how the hexagonal lattice could expand to form multiple pores. (E-H) Atomic scale models of a 6x6mer assembly with a central S1-lined pore. (E & F) Top and side views of the backbone. (G & H) Portions of cross-sections for aqueous and transmembrane domains with side-chains colored by element.

Large GM1-filled gangliosides Aβ42 hexamers may form larger hexamer-of-hexamers resembling those proposed for smaller hexamers (Fig. 14).

**Figure 14.**
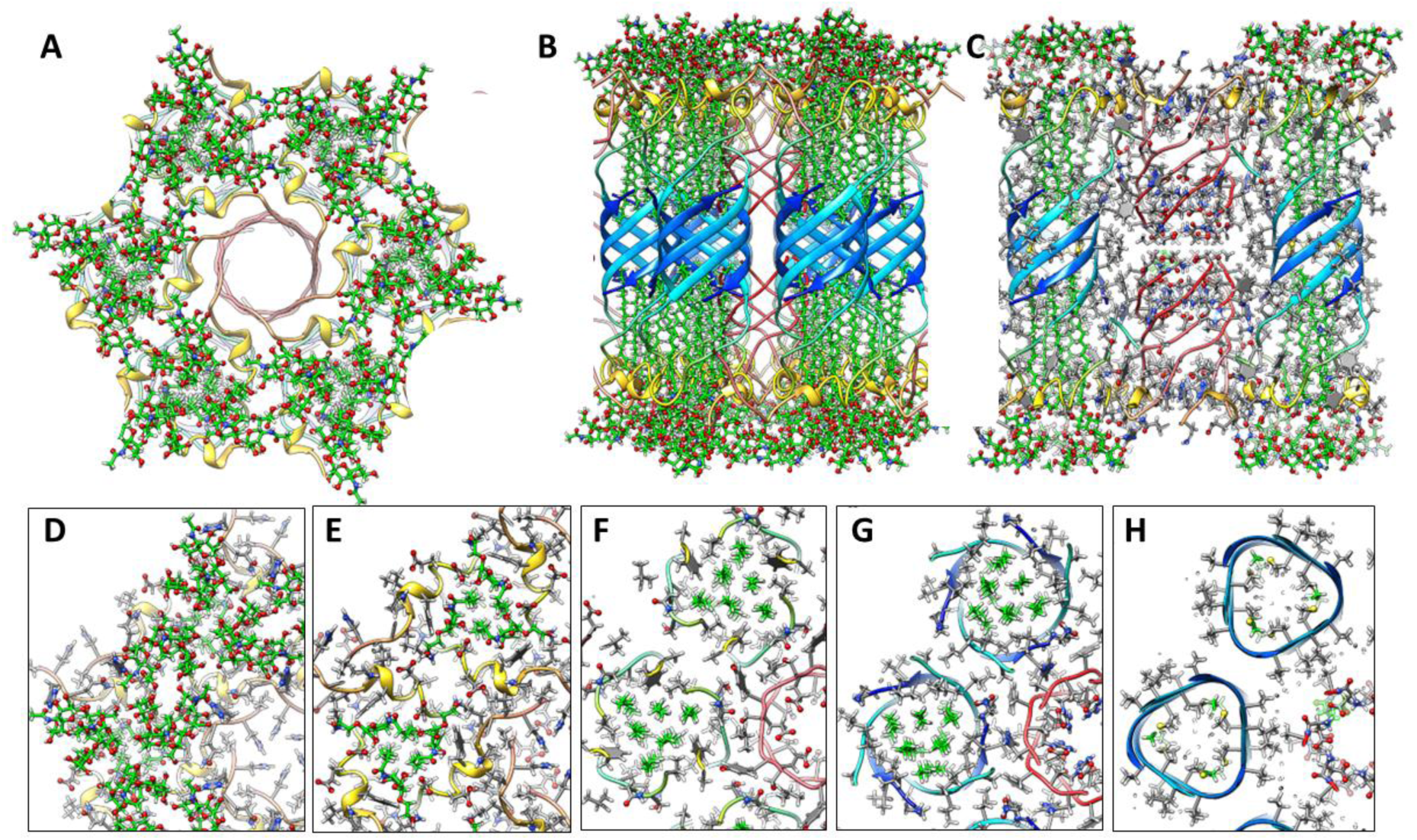
A hexamer-of-hexamers model composed of large 1Con2 hexamers in which S3 barrels are filled with GM1 alkyl chains. (A & B) Top and side views of backbone ribbon and GM1s. (C) Side cross-section showing S1 pore with side-chains. (D – H) Enlarged partial radial cross-sections with side-chains and GM1 (green carbons) colored by element centered between hexamers at regions of 2-fold symmetry beginning at the top and ending in the middle. Half of the S1 pore is on the lower right side in F-H.

### Dodecamer channel

The lining of the pore in the1Con2 dodecamer channel model of Fig. 15 is formed by two parallel 6-stranded S1 β-barrels that stack end on with N-termini Asp 1 residues meeting in the mid-region, much like the lining proposed above for the small 24mer channel and hexamer-of-hexamers model. S/N must be 2.0 for the S3 barrel to be large enough to surround the pore-lining S1 barrels. S2 segments form a ring of parallel α-helices in the aqueous phases surrounding the S1b portion of the S1 barrels, resembling how α-helices surround a parallel β-barrel in TIM α-β-barrel proteins (Wierenga, 2001). Strands of the outer S3 and inner S1 β-barrels are tilted by ∼55° and have S/N values of 2.0. Hydrophobic S1a side-chains (A2 and F4) interact with hydrophobic side-chains of S3, and those of S1b (Y10 and V12) interact with F19 of the S2 helix to form a cluster of aromatic side-chains (Fig. 15D). Electrostatic interactions are also favorable. Charged side-chains (D1, E3, R5, D7, E11 and H13) and the N-termini amine form salt-bridges that line the pore. Polar side-chains on the back side of S1 strands (H6 and S8) interact with the C-terminus carboxyl and polar side-chains at the end of S2 (D23) and linking S2 to S3. K16 of S2 salt-bridges to E22 of the adjacent helix and K28 at the beginning of S3 salt-bridges to the C-terminus carboxyl group of an adjacent S3 strand. Additional energetically favorable features include: tight side-chain packing between S1a and S2 barrels, burial or lipid exposure of hydrophobic side-chains and a high content of β and α secondary structure. Lithium ions khave been reported to inhibit onset of AD (Aron et al., 2025; Terao & Kodama, 2024). They were added in the negatively charged central region where they bind to Asp1 and Glu3 carboxyl groups.

**Figure 15.**
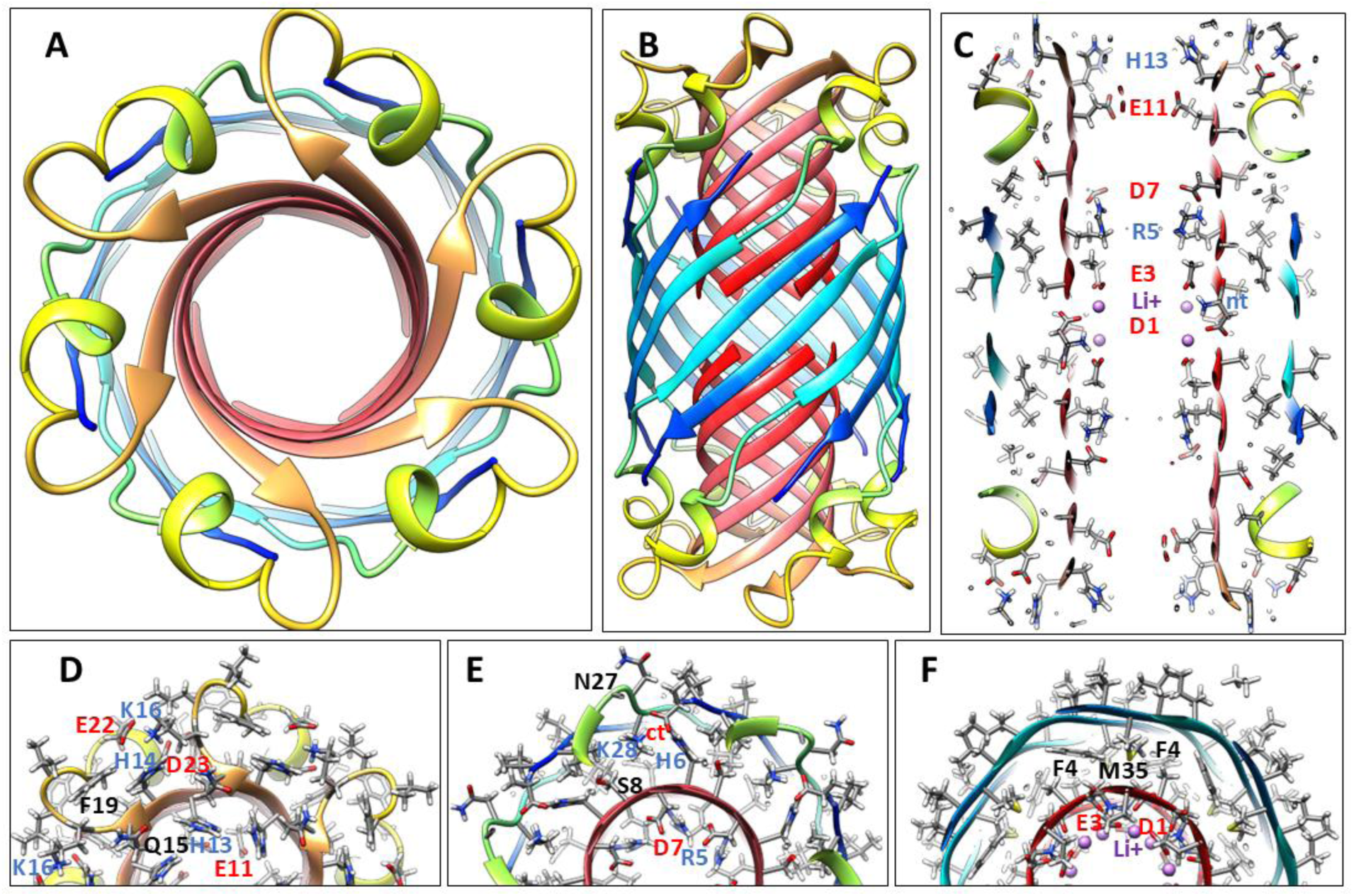
1Con2 model of a dodecamer channel with Li^+^ ions. (A & B) Top and side views of the ribbon backbone. (C) Side cross-section with side-chains. Charged residues lining the upper half of the pore and Li^+^ ions in near the central region are labeled. (D – F) Three cross-sections of the upper half of the channel with labeled side-chains for one subunit beginning at the top and ending in the center.

### 1Con2 model of an octadecamer channel

An octadecamer channel may form from insertion of three hexamers, analogous to the process illustrated in Fig. 10 for how four hexamers can form 24mer channels. The monomer folding motif and S3 strand tilt angles of the octadecamer model (Fig. 16) resemble that of the dodecamer; however, GM1 binds differently. An increase in the gap distance between the S1a and S2 barrels and the Glycine pleat provides enough space to accommodate GM1 alkyl chains between the S3 and S1 barrels in the Glycine pleat (Fig. 16 I). H14 side-chain binds to the GM1 carboxyl of the head-group. All charged moieties of the protein form salt-bridges. The pore through the nine-stranded parallel S1 β-barrels is naturally larger than that of the dodecamer.

**Figure 16.**
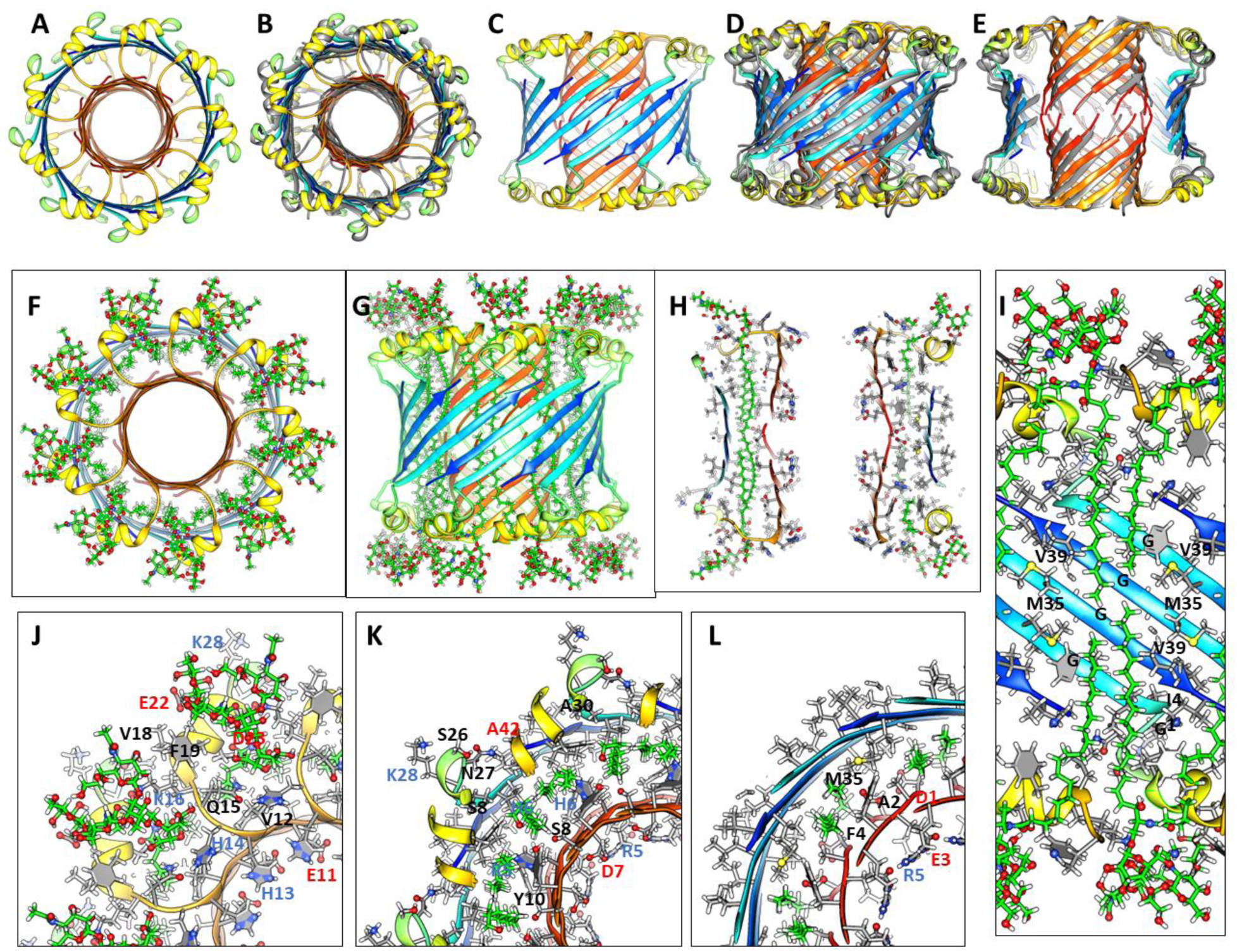
Atomic scale model of an Aβ42 1Con2 octadecamer with GM1 gangliosides bound between the outer S3 and inner S1 β-barrels. (A-E) Top (A & B) and side (C-E) views of the Aβ42 backbone before and after (superimposed in gray in B, D, and E) MD simulation for 200 ns. (E) A portion of the outer S3 barrel has been removed to reveal the S1 barrel. (F - H) Top and side view and cross-section of the backbone with GM1 ganglioside. (I) Enlarged side view cross-section of S2 helices and S3 strands viewed from inside the channel with side-chains colored by element and GM1 lipids. Glycines, M35 and V39 residues that interact with GM1 alkyl chains are labeled. (J - L) Portions of three radial cross-sections with GM1 and labeled side-chains beginning with the top surface and progressing to the center.

We use molecular dynamics to evaluate and refine our models. The octadecamer model simulated in a lipid bilayer is exceptionally stable; the secondary structure and position of the major segments are well maintained during a 200 ns simulation in a lipid bilayer (Fig.16 B, D & E) as are the position and conformations of the GM1 gangliosides.

### 1Con1 and 1Con2 36mer channels

Small and large 36mers of Figs. 17 and 18 are the largest channel models proposed to correspond to freeze-fracture images in Fig. 8. These may develop from the hexamer-of-hexamers construct of Fig. 14. Continuous S1 strands comprise 18-stranded parallel β-barrels with S/N = 1 for the S3 barrel of the smaller channel. GM1 gangliosides bind between the outer S3 and pore-lining S1 barrels; their head groups are located below the S2 helices.

**Figure 17.**
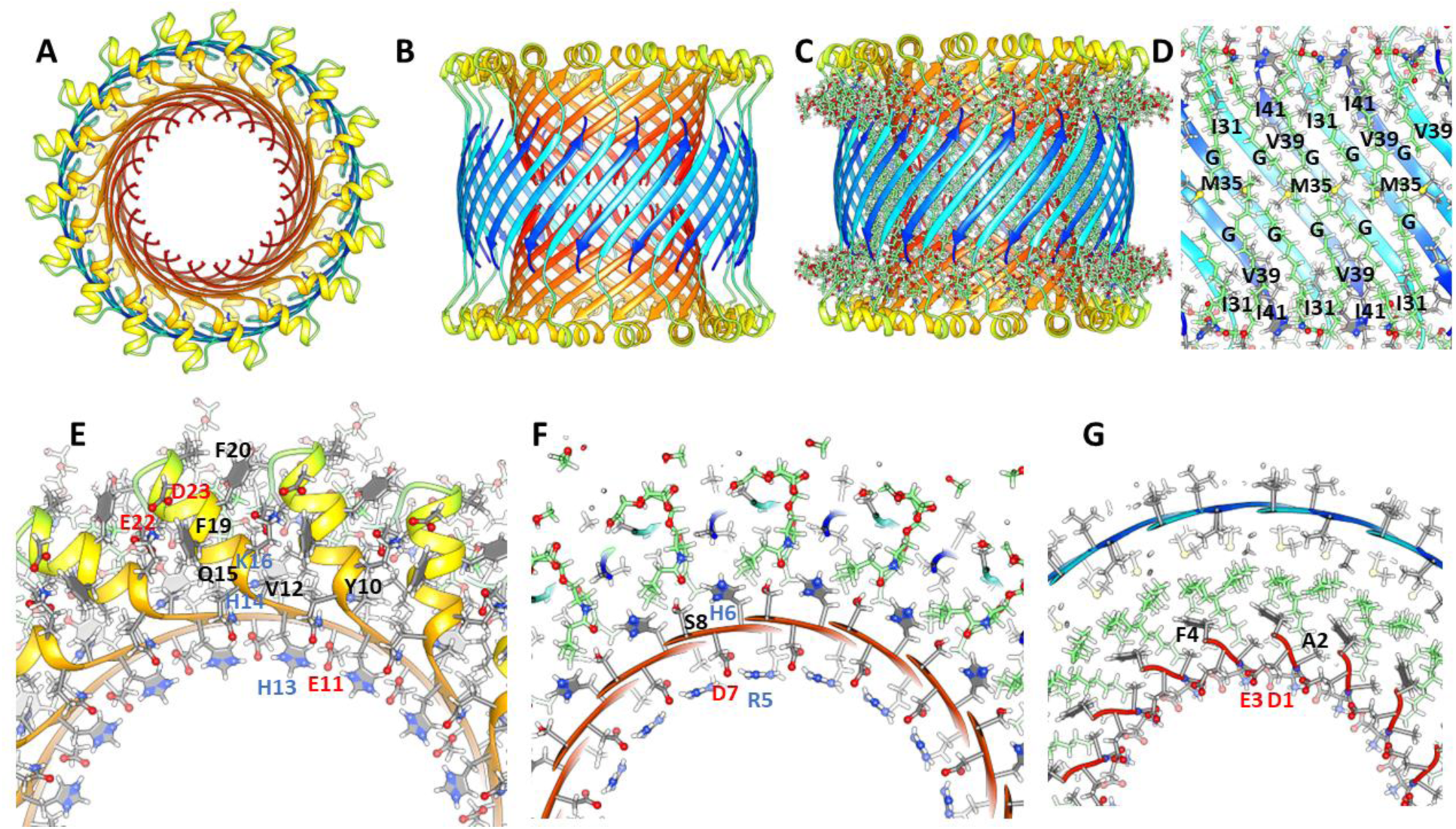
Atomic scale model of a small 1Con1 36mer-GM1 channel. (A-C) Top and side views of the backbone ribbon with GM1 gangliosides included in C. (D) Cross-section of the side-view showing how GM1 alkyl chains pack next to S3 side-chains. (E-G) Portions of radial cross-section with side-chains of the top, next lower, and middle regions.

**Figure 18.**
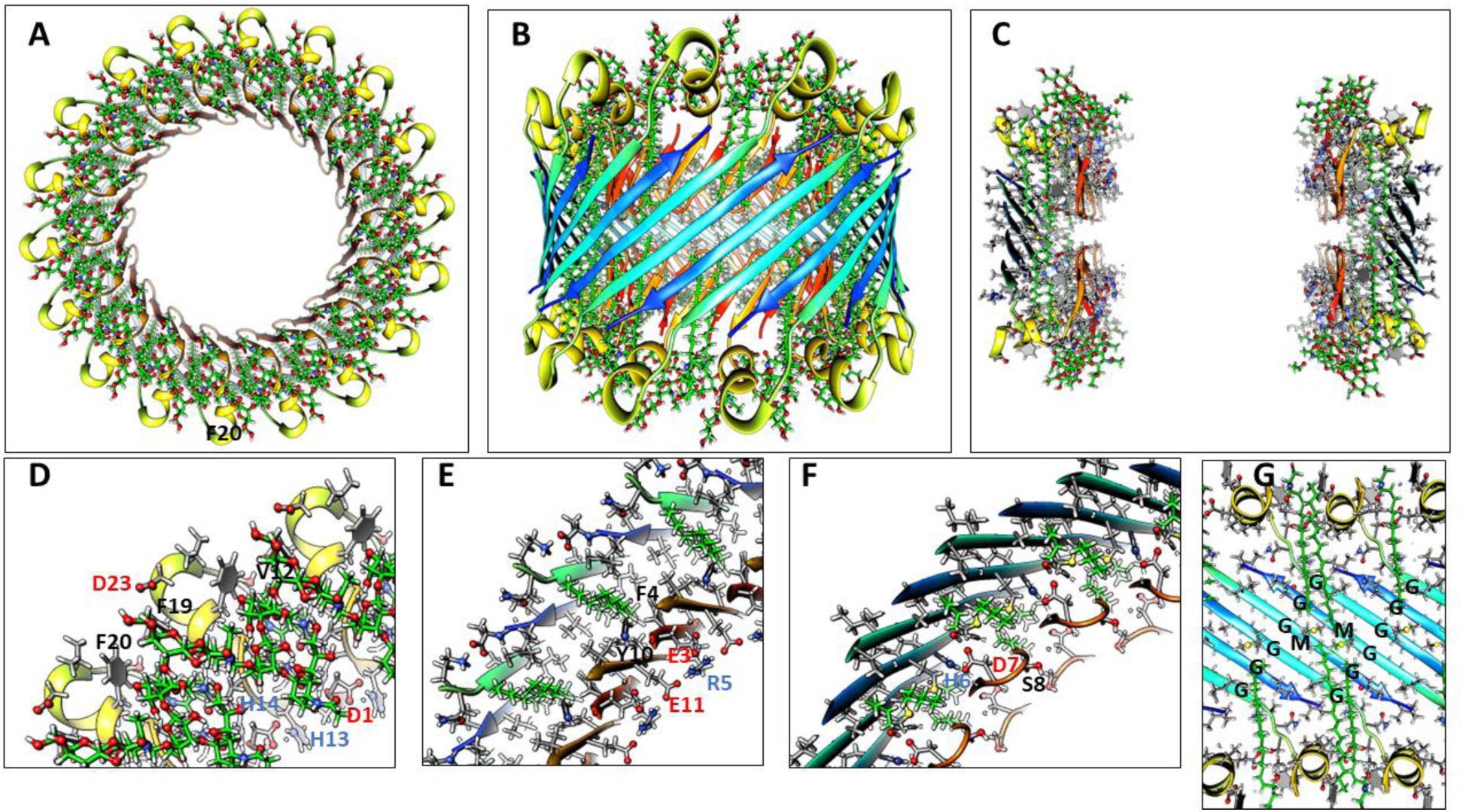
Atomic scale model of a large 1Con2 36mer-GM1 channel. (A & B) Top and side view with ribbon backbone and GM1 gangliosides. (C) Side view of a central cross-section with side-chains. (D-F) Partial radial cross-sections of the top, next lower, and middle layers. (G) Side cross-section showing how GM1 alkyl chains pack next to the S3 barrel’s interior wall.

The larger 1Con2 36mer model (Fig. 18) is shorter and wider than the 1Con1 model because the S3 strands are more tilted (S/N = 2, α = 55^◦^). S1a-S1b hairpins comprise two 36-stranded antiparallel pore-lining β-barrels with S/N = 1.0. GM1 headgroups are located between S2 helices. Their carboxyl group is surrounded by positive charges of His13, His14, Lys16 and the N-terminus amine. These are also counter-balanced by negatively charged carboxyl groups of

Asp1, Glu3, and Glu 11. Positively charged Arg5 side-chain interacts with Glu3 and Glu11 side-chains and the Asp7 and His6 side-chains interact in the hairpin turn.

### Plugged channels

FF images of Fig. 8 are proposed to be plugged channels. The atomic-scale model of Fig. 19 illustrates this type of structure for a 48mer in which a large 36mer channel of Fig. 18 surrounds a soluble dodecamer of Fig. 6.

**Figure 19.**
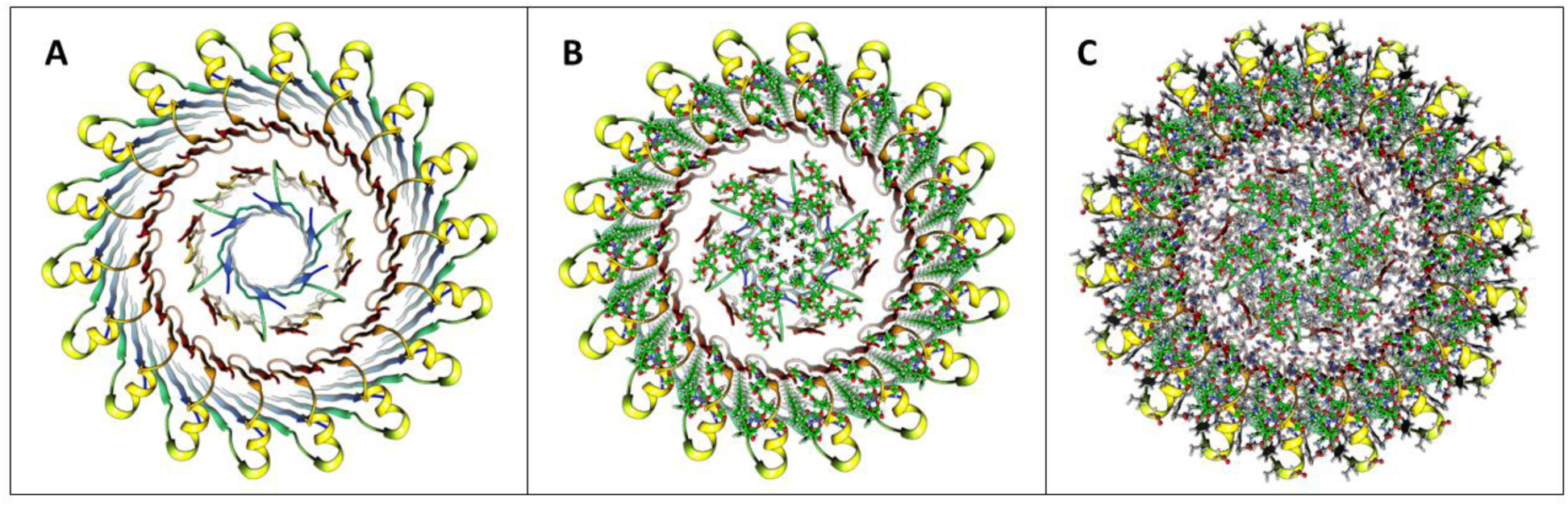
Model of a 48mer plugged channel in which a large 36mer channel surrounds a soluble dodecamer lipoprotein. (A) Backbone ribbon, (B) backbone with GM1, (C) backbone with side-chains and GM1.

### Atomic scale models of Aβ42 hexagonal lattices

Equilateral triangles are ideal for forming a honeycomb type hexagonal lattice (Fig 20). The S3 β-barrel of our large hexamer model is filled with GM1 alkyl chains. Parallel 6-stranded S1 β-barrels (red strands) form small pores between six hexamers in our lattice model.

**Figure 20.**
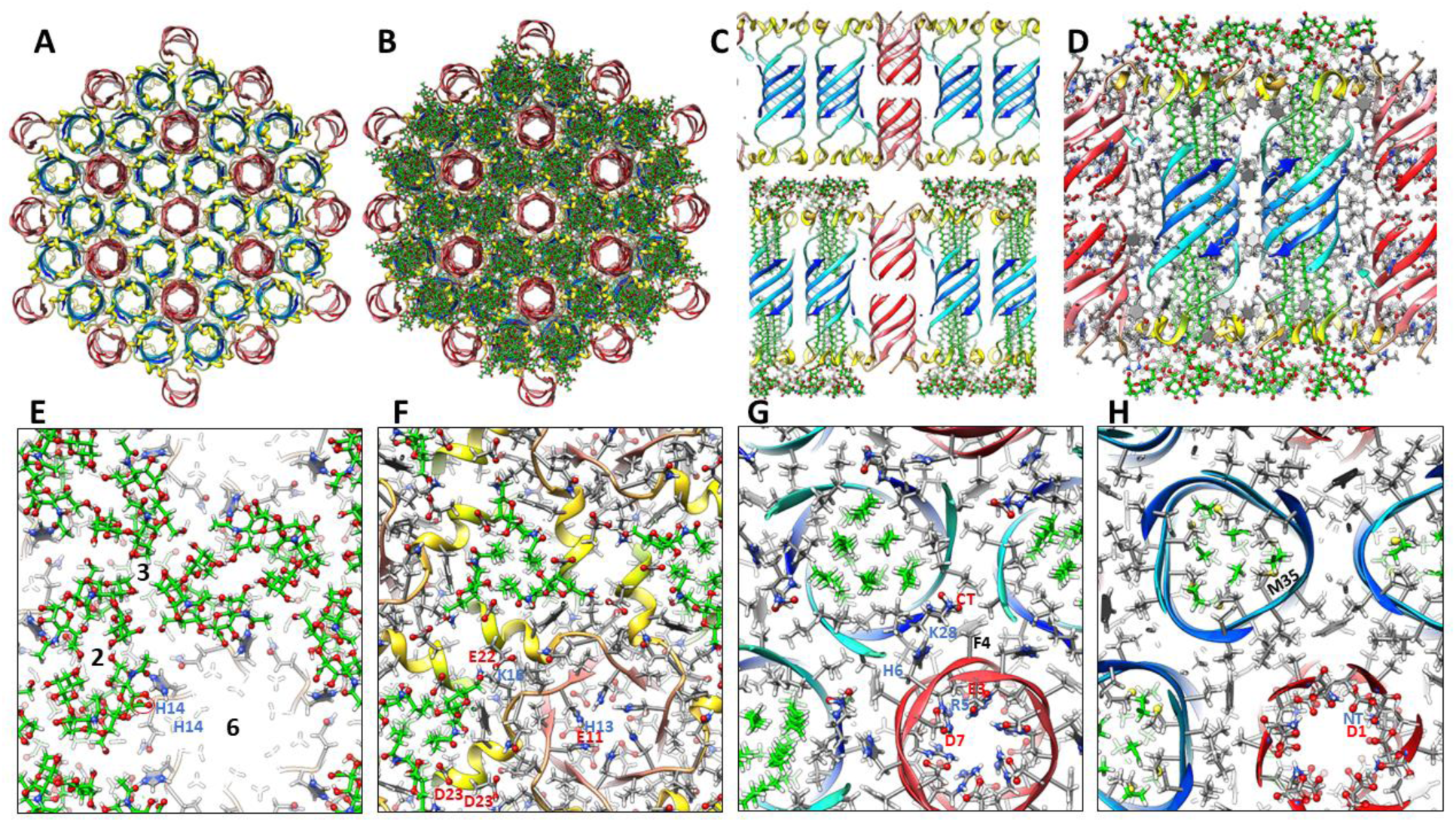
Model of Aβ42 large hexamer channel lattice with associated GM1. (A &B) Top view of ribbon backbone alone and with GM1s. Six hexamers contribute to 6-stranded parallel S1 β-barrels of the red-colored pores. (C) Side views for a portion of the central row with backbone ribbons alone (top), with GM1s (bottom), and (D) with side-chains and GM1s (green carbons) colored by element. (E-H) Enlarged radial cross-sections of regions with 2-fold, 3-fold, and 6-fold symmetries beginning at the top and ending in the middle. (E) The top layer GM1 headgroups; axes of 2-,3-, and 6-fold symmetries are numbered. His14 side-chain binds to GM1 carboxyl group. (F) The next layer shows S2 α-helices (yellow) and GM1 headgroups; His13 binds to Glu11 and Lys16 binds to Glu22. (G) Green GM1 alkyl chains are inside the S3 barrel, Lys28 binds to the C-terminus carboxyl, and Arg5 binds to Glu3 and Glu11. (H) Asp1 carboxyl and N-terminus amine of the two S1 β-barrels form salt-bridges in the mid-region and Met 35 side-chains occupy the space between ends of the inner and outer GM1 alkyl chains.

The 6-fold radial symmetry of dodecamer TMO and channel models may allow them to aggregate into triangular-type hexagonal lattices. S3 barrels of the TMO dodecamer model are filled with GM1 and cholesterol lipids and the GM1 carboxyl group binds to His13 and His14 side-chains (Fig. 21). These dodecamers pack tightly together in a hexagonal lattice without lipids between dodecamers and S1 β-hairpins form a 4-stranded β-sheet connecting dodecamers with their numerous charged side-chains expose to the aqueous phase and forming salt-bridges.

**Figure 21.**
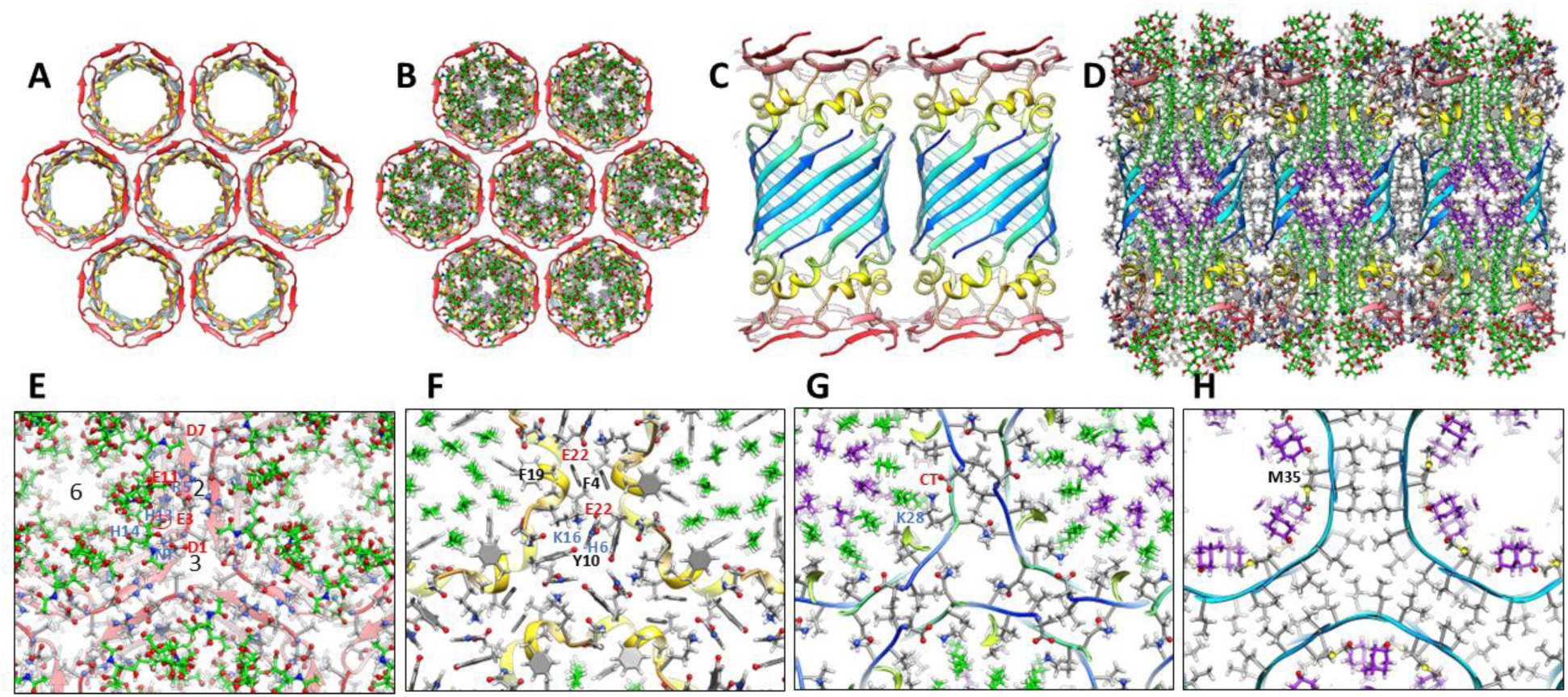
A hexagonal lattice model of seven dodecamer TMOs. (A) Top view of a central Aβ42 dodecamer TMO backbone ribbon surrounded by six other dodecamers. (B) GM1s (green carbons) and cholesterols (purple carbons) are bound inside the dodecamers. (C) Side view of two dodecamer backbone ribbons. (D) Side view cross-section of the central row including side-chains colored by element and GM1s and cholesterol (purple). (E - H) Enlarged radial cross-sections for a portion of the lattice that includes regions of 2-, 3-, and 6-fold symmetries and GM1. (E) The top cross-section includes the S1 β-hairpins and GM1 headgroups. The carboxyl group of one GM1 is circled in red and symmetry axes are labeled. (F) Next layer down showing S2 α-helices and top portions of GM1 alkyl chains. (G) Third cross-section with top portions of the S3 β-barrel, bottom portions of GM1 alkyl chains, and cholesterols. (H) Middle cross-sections showing Met35 side-chains, interactions with ends of cholesterol inside each S3 barrel and interactions among exterior S3 alkyl chains between dodecamers.

Our dodecamer channel models may form a similar hexagonal lattice (Fig. 22). In this model GM1 lipids bind between dodecamers at regions of 3-fold symmetry.

**Figure 22.**
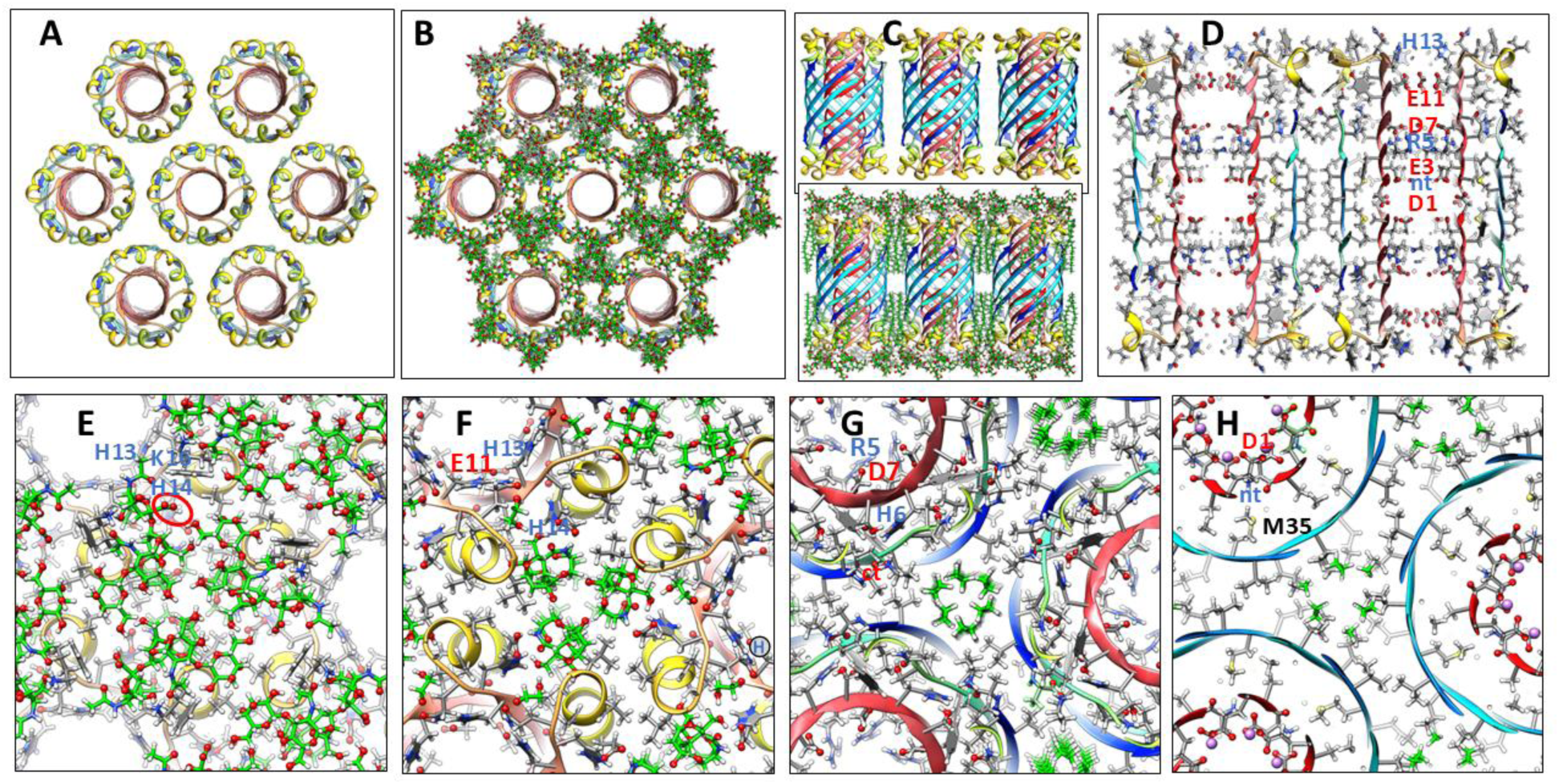
Hexagonal lattice model of dodecamer channels. (A) Ribbon backbone of seven 12mer channels viewed from the top. (B) Same as (A) with 24 GM1s (green carbons) added in the regions of 3-fold symmetry between 24mers. (C) Side view of three dodecamer backbones without and with GM1s. (D) Cross-section side view showing interactions of side-chains between two dodecamers at a region of 2-fold symmetry. Charged side-chains of half of the S1-lined pore are labeled. (E-H) Enlarged portion of lattice showing regions of 2-, 3-, and 6-fold symmetries. (E) Top layer showing GM1 headgroups with GM1 carboxyl circled in red and interactions with His13, His14, and Lys16. (F) Next layer down showing S2 helices and GM1 headgroups with some charged side-chains labeled. (G) Third cross-section showing beginnings and ends of S3 β-strands (blue) and S1 pore lining (red). (H) Mid-region cross-section showing ends of GM1 alkyl chains and interactions of the N-termini of S1 pore-lining barrels.

The 12-fold radial symmetry of our 24mer channel models may also allow them to aggregate into a hexagonal lattice. A lattice of the smaller 1Con1 models resembles that of Fig. 23 in which GM1 gangliosides bind in regions of 3-fold symmetry between channels and S1 segments are single β-strands (not illustrated). In contrast, GM1s bind between the S1 and S3 barrels in the larger 1Con2 models and additional GM1s or other lipids would occupy some of the space between the channels (Fig. 23).

**Figure 23.**
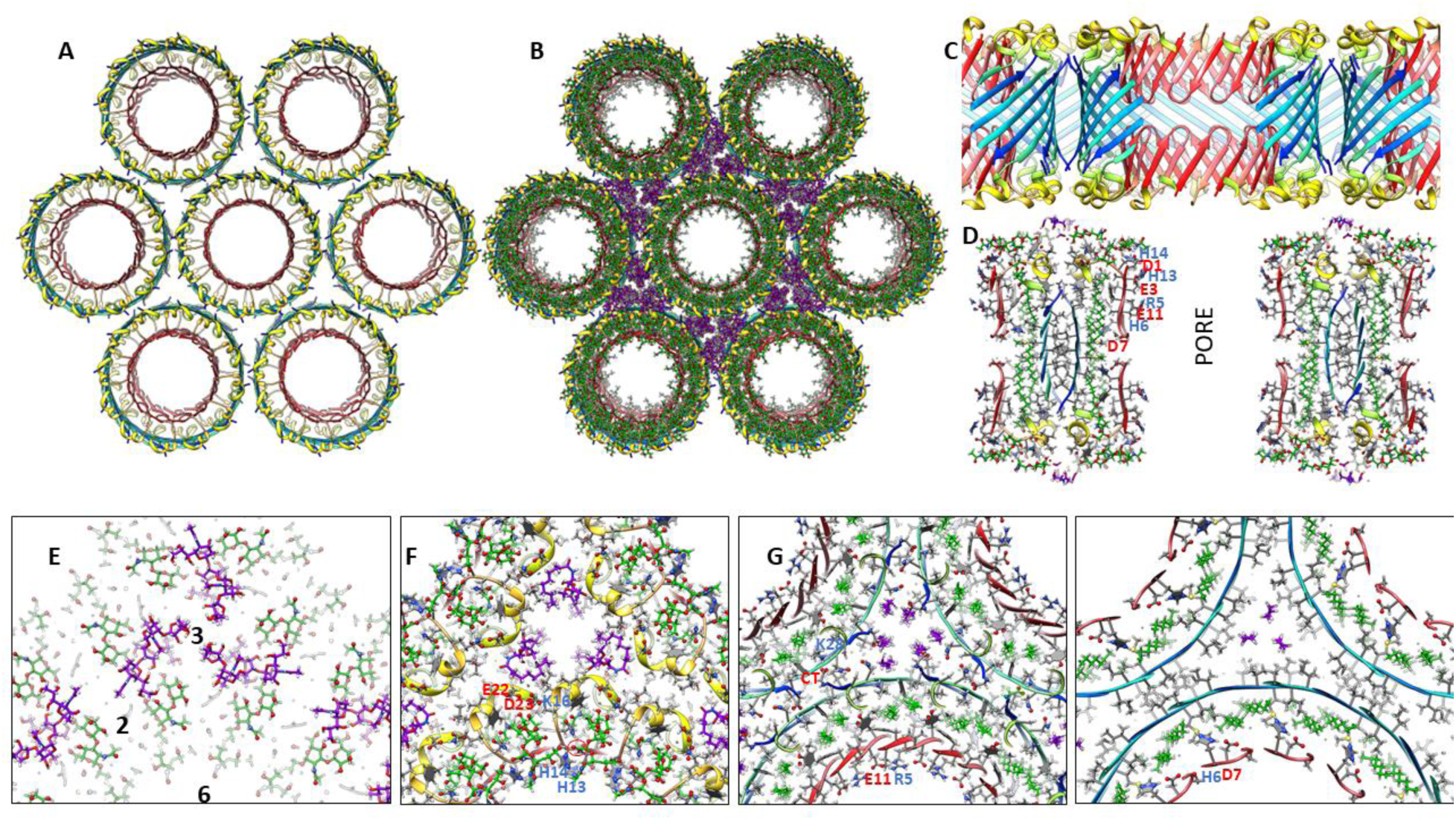
Hexagonal lattice model of large 24mer channels. (A) Ribbon backbone of seven 24mer channels viewed from the top. (B) Same as (A) with 24 GM1s (green carbons) added between the S3 and S1 β-barrels and six GM1s (purple) in regions of 3-fold symmetry between 24mers. (C) Side view cross-section of the backbone for the central region that includes the S1 pore. (D) Thin cross-section with side-chains and GM1s. Charged side-chains of the S1-lined pore are labeled. (E-H) Enlarge portion of lattice showing regions of 2-, 3-, and 6-fold symmetries. (E) Top layer showing GM1 headgroups with symmetry axes labeled. (F) Next layer down showing S2 helices and GM1 headgroups with GM1 carboxyl circled in red and some charged side-chains labeled. (G) Third cross-section showing beginnings and ends of S3 β-strands (blue) and S1 β-hairpins (red). (H) Mid-region cross-section showing ends of GM1 alkyl chains and cholesterols and turn region of S1 β-hairpins. Notable charged side-chains are labeled.

### Hexagonal Lattice of 1Con Channels

The above hexagonal lattice models suggest an intriguing possibility: perhaps the different lattices should not be considered independently but rather as different conformational states of the same assembly; increases in lateral membrane tension may stimulate transitions from closed or smaller channels to fewer but larger channels. Thus, a relatively large lattice could be a mechanosensitive complex that transitions from closed to large pores. Such complexes may form when membranes fuse; *e.g.*, when fusion pores release neurotransmitter at synapses. Fig. 24 illustrates the basic concept. Each TMO and channel assembly is composed of 288 Aβ42 peptides. S/N is 1.0 for S3 barrels of the top row. S3 barrels of the bottom row are shorter and wider due to their S/N values of 2.0. This type of transition from tall/thin/closed channels to short/wide/open channels has been reported for other mechanosensitive channels such as MscL (Sukharev et al., 2001).

**Figure 24.**
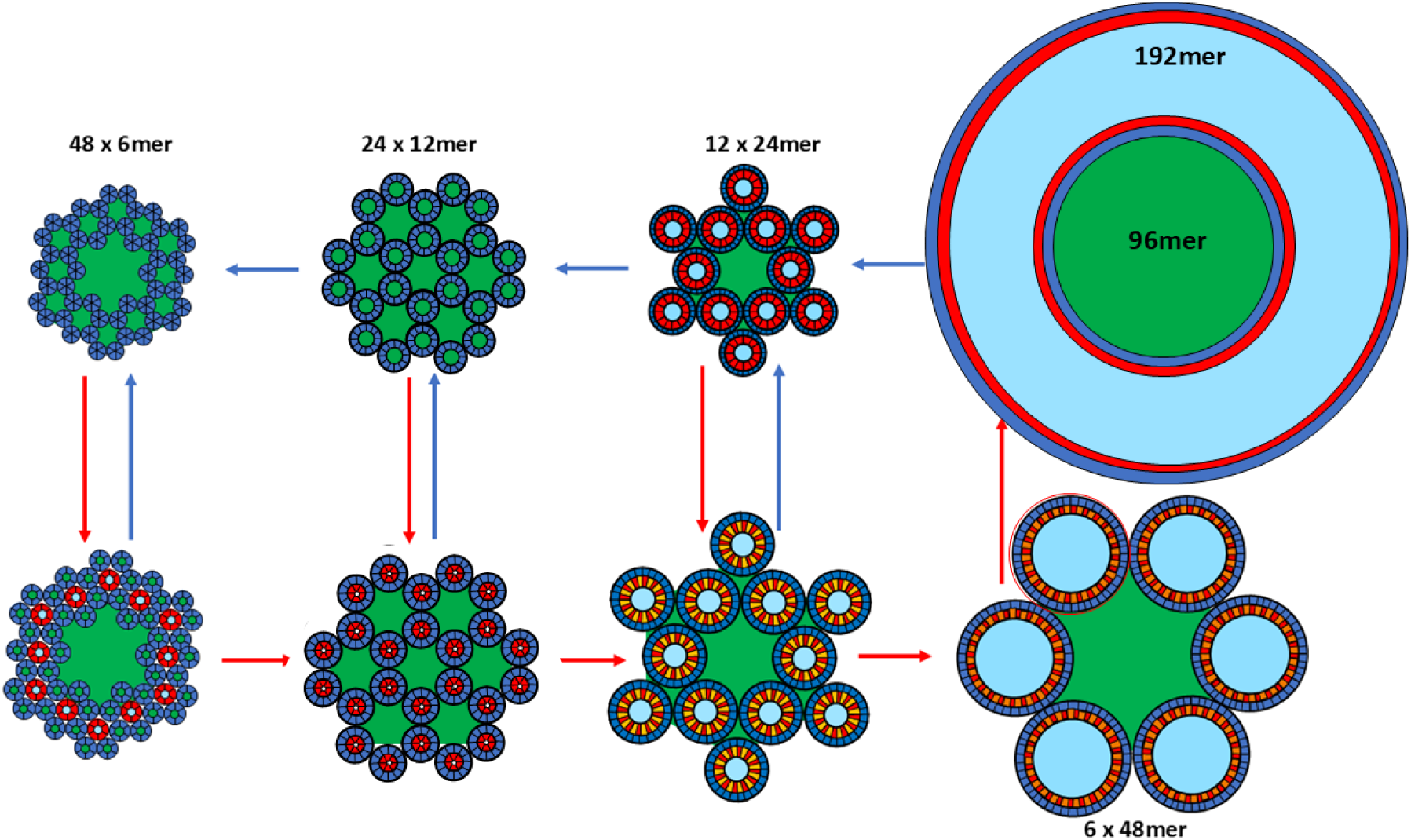
Model of mechanosensitive gating of an Aβ42 hexagonal lattice. Each of the eight schematics has 288 1Con type Aβ42 peptides. Lipid- and water-filled regions are green and light blue. S3 barrels have S/N values of 1.0 (α = 36^⸰^) for the top row and 2.0 (α = 55^⸰^) for the bottom row. Pore sizes increase from left to right and from top to bottom except for the plugged-pore model in the top right. Transitions from left to right occur when two smaller TMOs or channels merge to form a channel with twice as many peptides.

However, this model has a second, more drastic type of mechanosensitivity. In this example, TMOs of a lattice comprised of 48 hexamers can merge to create a lattice of 24 dodecamers with small pores; the dodecamers may then merge to form twelve 24mers with medium sized pores; 24mers may merge to form six 48mers with very large pores. The final proposed step occurs by the mechanism proposed above for formation of channels surrounding a lipoprotein type assembly: *i.e.*, the 48mers split with the third of the subunits facing inwardly and combining to form a central lipoprotein with 96 subunits, and the two-thirds facing outwardly combining to form the surrounding 192mer channel.

### Atomic Force Microscopy of Aβ42 Assemblies in Supported Bilayers

Mrdenovic et al., (2019) (Mrdenovic et al., 2019) used atomic force microscopy imaging to study effects of both large and small Aβ42 oligomers on supported bilayers composed of brain total lipid extract. They reported that small Aβ oligomers created large pores through the bilayer with a diameter of ∼ 44 nm and soluble lipoprotein discs with diameters ranging from ∼ 20 to 40 nm (Fig. 25 A-D), and that the membrane eventually disintegrates into lipoprotein clusters. We suggest that these huge holes may develop from a hexagonal lattice of nineteen 24mers plus some surrounding peptides (Fig. 25 E). In this illustration each 24mer of the outer ring of twelve 24mers splits and merges to form a 168mer channel and a l20mer soluble β-barrel that surrounds the core of seven 24mers. Once the blocked channel (Fig. 25F) is formed, the lipoprotein of the core exits into the aqueous phase to form the lipoproteins of Fig. 25 D, leaving behind the large hole of Fig. 25 A & B. Eventually the β-barrels forming the wall of the hole collapse into its constituents to form a randomized glob of TMOs that could range in size from 4 to 32 subunits, corresponding to the Fig. 25 C image. This speculative process would probably require higher concentrations of Aβ42 than exist in the brain, but if it does occur even rarely, it could be deadly for neurons.

**Figure 25.**
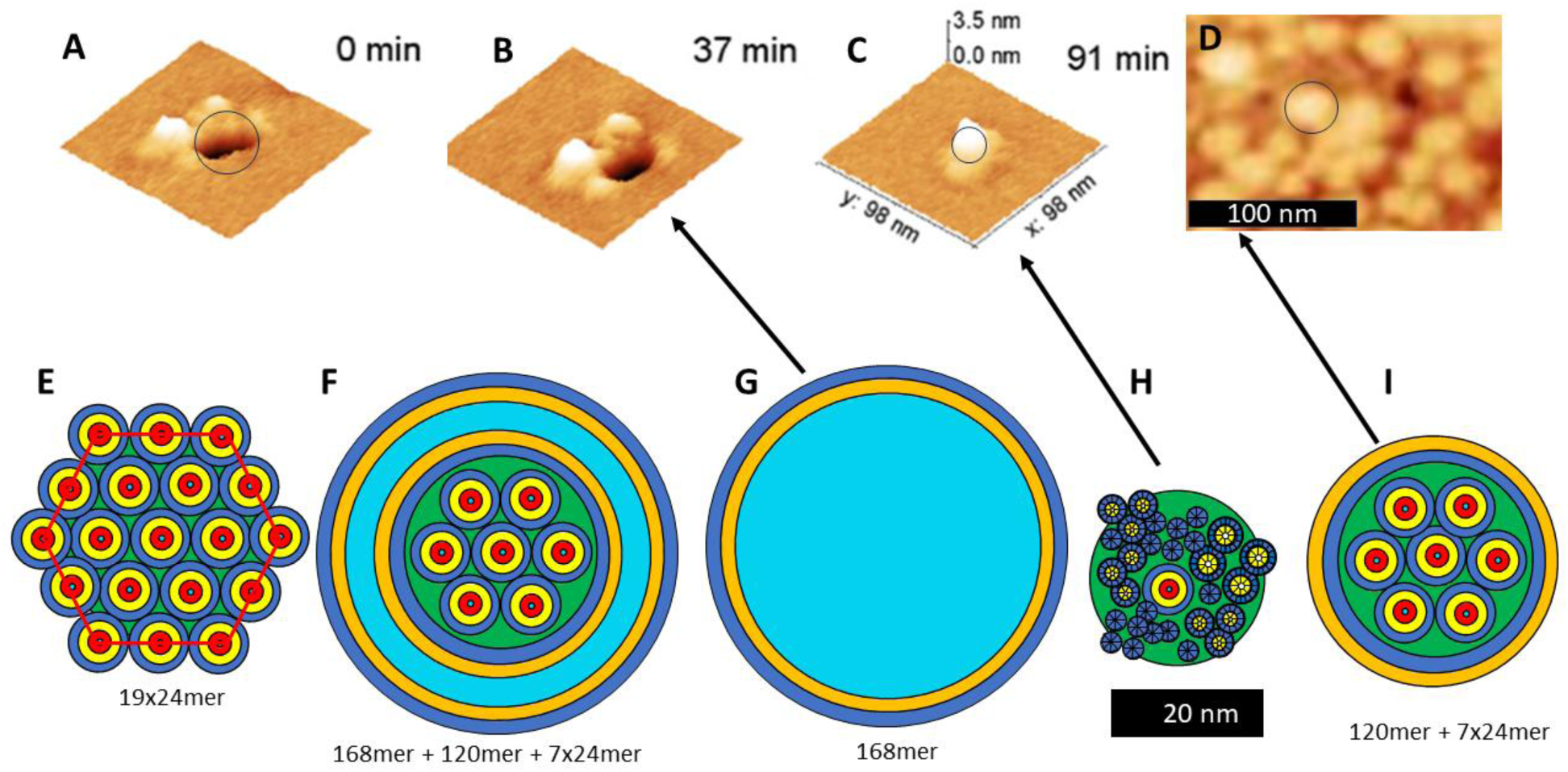
AFM images of large transmembrane pores and soluble lipoprotein plus schematic models of Aβ42 structure types possibly associated with these structures. AFM images of Aβ42 assemblies (A-C) in supported bilayers and (D) soluble lipoprotein (adapted from Mrdenovic et al., (2019) (Mrdenovic et al., 2019)). (E) Schematic of a hexagonal array of nineteen 24mer Aβ42 channels. (F) Each 24mer of the outer ring of twelve 24mers splits and merges with 2/3rds of the peptides forming a giant channel and 1/3rd forming a lipoprotein with the core seven 24mers in its center. (G & H) The lipoprotein assembly enters the aqueous phase leaving behind a giant channel that corresponds to images of (A) and (B). (I) The β-barrel wall of the giant channel collapses into a glob of smaller components corresponding to the image of (C).

## Conclusions and Discussion

Decades of AD research have not produced cures or even effective inexpensive treatments. This lack of progress may be related to our poor understanding about molecular, cellular, and biological mechanisms responsible for the gradual deterioration characteristic of AD. There is little doubt that Aβ is involved. Most efforts regarding Aβ have targeted fibrils that are major components of amyloid plaques. But plaques become most prevalent in late stages of AD after much damage has already occurred. Antibody treatments that remove only plaques have proven relatively infective, often toxic, and expensive; whereas those that also target Aβ oligomers slow the disease (Terao & Kodama, 2024).

Earlier intervention during or prior to the Mild Cognitive Impairment (MCI) stage may hold more promise. One of the earliest detectable symptoms of the onset of AD is an increase in the ratio of Aβ42 to Aβ40. Relatively small Aβ42 oligomers are more toxic than any other known form of Aβ (large oligomers, fibrils, or other common variants) and are the only Aβ assemblies that have been demonstrated to form ion channels in neuronal plasma membranes (Bodie, 2017).

In this and the accompanying paper we postulate that Aβ42 can adopt relatively simple concentric β-barrel motifs consistent with the following types of assemblies: relatively small soluble oligomers composed of 4, 6, 8, 12, 16, and 18 subunits, beaded annular protofibrils in which each bead is an oligomer and the number of oligomers vary, smooth annular protofibrils that develop from mergers of the beads, membrane surface assemblies that form after soluble oligomers bind to membranes, transmembrane oligomers that result from the reorientation of surface assemblies, small transmembrane channels that develop from mergers of TMO assemblies, larger transmembrane channels and possibly soluble lipoproteins that develop from mergers of smaller channels or larger TMO assemblies. Our models are consistent with low resolution microscopy images of all these assembly types and with other experimental findings as well as molecular modeling criteria.

Making our structural models consistent with most experimental results forced us to develop a multitude of models, leading to the conclusion that Aβ42 assemblies are highly polymorphic. Extreme polymorphism may explain why distinct structures have been so difficult to determine. Polymorphism also forced us to search for a relatively simple protein folding motif applicable to all the polymorphs. The common feature of all our current models is that S3 strands comprise symmetric antiparallel β-barrels with both radial and P2 symmetries. Sizes of these barrels depend on the number of subunits in the assembly and tilts of the S3 strands. For 1Con models in which all subunits have the same conformation, the tilts are constrained to 36^⸰^ or 55^⸰^. S3 barrels comprise a hydrophobic core for soluble oligomers and an outer lipid-exposed barrel for transmembrane models. Outer S1 and/or S2 β-barrels shield the exterior of S3 barrels of soluble oligomers from water; whereas, S1 β-barrels shield the interior of transmembrane channel S3 barrels from water by lining the pore.

The calcium hypothesis of AD proposes that Ca^2+^ homeostasis, essential to normal neuronal functioning, becomes disrupted by AD (see Wang et al 2025 for recent review). The early finding that Aβ channels are permeant to Ca^2+^ (Arispe et al, 1993) has been verified in numerous laboratories (Li et al., 2023). S1 is a highly charged segment with negatively charged residues at positions 1, 3, 7, and 11. Our hypothesis that S1 lines the pore of 1Con type channels is thus consistent with Ca^2+^ permeation through both plasma and organelle (e.g. mitochondria) membranes. In contrast, Li^+^ ions may inhibit onset of both Mild Cognitive Impairment (MCI) and AD (Aron et al., 2025; Terao & Kodama, 2024). Li^+^ binding sites typically involve multiple carboxyl groups from Asp and/or Glu side-chains (Dutta et al., 2014). These types of sites could occur in our models between the Asp1 and Glu3 side-chains in the central region of the pore where the N-termini of the two putative S1 barrels meet (Fig. 15). The Lithium cations remained bound to these groups throughout relatively long MD simulations. If Li^+^ binds tightly to these sites, its presence would make this portion of the pore less electronegative and thus make the pore less permeant to Ca^2+^. Aβ channels appear to be inhibited by the larger cations Zn^2+^ and Al^3+^ (Kagan et al., 2002) that may also interact with these sites. Permeation of Ca^2+^ through Aβ42 channels of plasma and/or organelle (e.g. mitochondria and Ca^2+^ stores) membranes could disrupt calcium regulation and kill cells.

Aβ42 channels are unlikely to act alone. GM1 gangliosides and cholesterol affect their toxicity and ability to form channels. Thus, it is important to include them in molecular models and to consider inhibition of their binding to the channels as a possible treatment (Fantini et al., 2020). Most of our soluble oligomer models include GM1, consistent with the lipid chaperone hypothesis (Sciacca et al., 2020).

The possibility that some Aβ42 channels are functionally vital may complicate treatments that target them. For example, dodecamer Aβ42 channels could be functional but become pathogenic if they aggregate into large assemblies or morph into larger channels. These transitions should become more probable as the concentration of Aβ42 increases. Thus, strategies to inhibit these transitions should be considered. If AD is caused by an overabundance of Aβ42, then it may be necessary to regulate their concentration rather than to completely inhibit them.

An intriguing possibility suggested by our models is that Aβ42 channels have two mechanisms for being mechanosensitive. The first is that lateral tension could favor transitions of the small-tall channels (S/N = 1.0) to larger-shorter channels (S/N = 2.0). The second is that lateral tension could stimulate mergers of smaller assemblies (TMOs or small channels) to form larger channels. To our knowledge, this hypothesis for mechanosensitivity has not been tested.

Several features of our current models are not new. Our proposals that channels have both radial and P2 symmetries, that some channels have twelve identical subunits, that S2 segments form surface α-helices, and that S1 segments line the pore were made three decades ago ((Durell et al., 1994), see supplement). Our 2010 models (Shafrir, Durell, Arispe, et al., 2010; Shafrir, Durell, Anishkin, et al., 2010; Yun et al., 2011) postulated that S3 forms antiparallel β-barrels that are concentric with S1 and/or S2 β-barrels and utilized β-barrel theory to constrain the models. The major additions in our current models are incorporation of GM1 lipids into the structures, proposing that TMO assemblies may morph into channels, increasing the plausible number of transmembrane Aβ42 models, proposing channels may aggregate into 2-D lattices that may be mechanosensitive, and proposing that some large channels are partially plugged by soluble lipoproteins that may then exit to the aqueous phase.

Our new and radical departure from conventional thinking should be considered skeptically. Even if basic concepts are correct, fine details of the atomic-scale models are sure to have errors and illustrations of transitions are likely over simplified. Better and higher resolution data are urgently needed. The conclusions section of the accompanying paper suggests ways our models can be tested and how higher resolution structures might be obtained.

## Acknowledgements

This work was supported in part by the Intramural Program of the National Institutes of Health, National Cancer Institute, Center for Cancer Research.

