## Supplemental 2 Figures for "The concentric β-barrel hypothesis for amyloids: Models of soluble and transmembrane amyloid-β42 oligomers and channels composed of identical subunits and GM1 gangliosides"

*Contact information:

H. Robert Guy


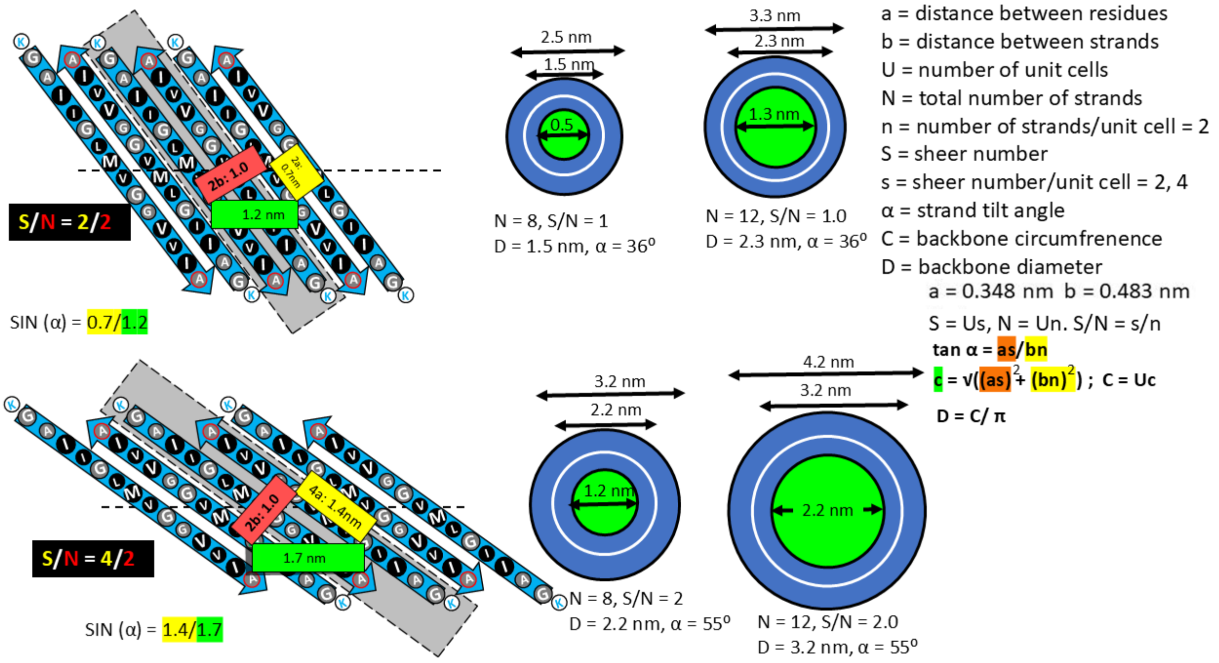


Figure S1. Beta barrel theory and definition of terms for 1Con models. The central unit cell in the flattened schematics has a gray background.

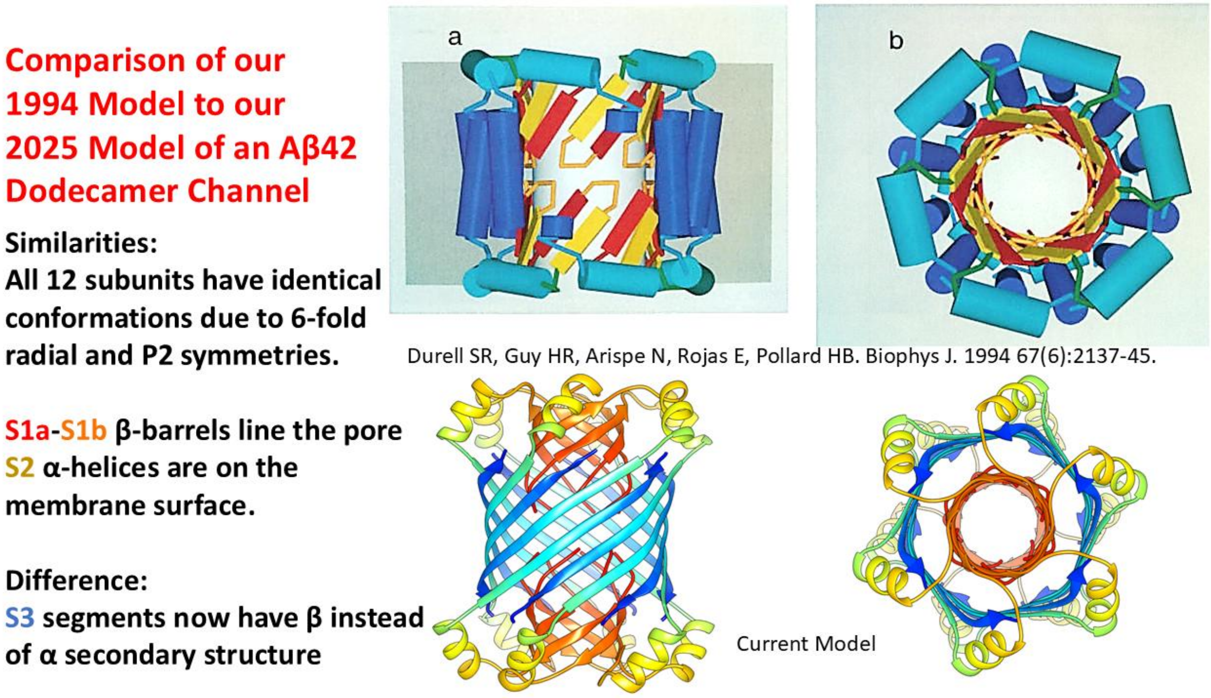


Figure S2. Comparison to first and current Aβ42 dodecamer channel models. (a & b) Top and side views of backbone. S2 helix is cyan in the ’94 model.
